# CRISPR/Cas9 and FLP-FRT mediated multi-modular engineering of the *cis*-regulatory landscape of the bithorax complex of *Drosophila melanogaster*

**DOI:** 10.1101/2022.06.12.495802

**Authors:** Nikhil Hajirnis, Shubhanshu Pandey, Rakesh K Mishra

**Author notes:** For correspondence –. Department of Anatomy and Neurobiology, University of Maryland, Baltimore, United States of America (Nikhil Hajirnis).

## Abstract

The Homeotic genes or *Hox* define the anterior-posterior (AP) body axis formation in bilaterians and are often present on the chromosome in an order which is collinear to their function across the AP axis. The expression pattern of *Hox* genes is attributed to the *cis*-regulatory modules (CRMs) that regulate the genes in a segment-specific manner. In the bithorax complex (BX-C), one of the two *Hox* complexes in *Drosophila melanogaster*, even the CRMs consisting of enhancers, initiators, insulators, and Polycomb/trithorax response elements are organized in order that is collinear to their function in the thoracic and abdominal region. Much of these findings are based on the analysis of hundreds of mutations in the BX-C. However, targeted genomic rearrangements comprising of duplications, inversions, etc., that can reveal the basis of collinearity and the number of regulatory modules with respect to body segments have not been reported. In the present study, we generated a series of transgenic lines with the insertion of FRT near the regulatory domain boundaries, to shuffle the CRMs associated with the posterior *Hox*, *Abd-B*, of the BX-C. Using these FRT lines, we created several alterations such as deletion, duplication, or inversion of multiple CRMs to comprehend their peculiar genomic arrangement and numbers in the BX-C.

## Introduction

Eukaryotic gene regulation is a complex process facilitated by a combination of *cis*-regulatory elements (CREs) and *trans*-acting factors (TAFs). The fine-tuning of expression of a gene is performed by several *cis*-regulatory elements that include initiator elements, enhancers, repressors, Polycomb/trithorax response elements and promoter targeting sequences (Narlikar and Ovcharenko 2009; Crocker et al. 2016; Iampietro et al. 2010; Gaston and Jayaraman 2003; Zhou and Levine 1999; Ringrose and Paro 2004). In *Drosophila melanogaster* genome, these elements are often populated in 15-20 Kb regions that regulate the activity of associated genes. Such regions where several CREs populate together and are responsible for the expression of a gene in a cell, tissue, or segment-specific manner are called *cis*-regulatory modules (CRMs) (Ho et al. 2009; Starr et al. 2011; Bekiaris et al. 2018). Plentiful studies have been dedicated towards understanding the role of each of these elements individually (Polychronidou and Lohmann 2013; Kyrchanova et al. 2011; Cléard et al. 2006; Akbari et al. 2008; Narlikar and Ovcharenko 2009). However, only a handful of them addressed the collective functioning of the CRMs (Lelli et al. 2012; Zinzen et al. 2009; Starr et al. 2011; Hajirnis and Mishra 2021). One such region in the *Drosophila* genome that heavily relies on the collective functioning of the CRMs, includes a set of developmentally important genes, called *Hox*. These genes code for transcription factors that bind to the DNA in a sequence-specific manner. The expression pattern of these genes defines the anterior-posterior body axis of a developing bilaterian embryo (Foronda et al. 2009; Hajirnis and Mishra 2021).

Homeotic genes or *Hox* were discovered in *Drosophila melanogaster* wherein they are present in a spatially colinear manner on the chromosomes (Lewis 1978). That is, the genes are arranged in a complex and, the genes that are present at one end of the complex are responsible for the development of the anterior part of an embryo while the genes present at the other end are responsible for the development of the posterior segments. The *Drosophila Hox* genes are arranged in two clusters, the Antennapedia complex (ANT-C) and the bithorax complex (BX-C) (Kaufman et al. 1990; Peifer et al. 1987).

The posterior-most *Hox* gene, *Abd-B* is regulated by four distinct CRMs, *infradbominal-5* (*iab5*), *iab6*, *iab7,* and *iab8/9*. Notably, in addition to the arrangement of *Hox*, even the CRMs of the BX-C genes are spatially arranged in a manner colinear to the A-P body axis that they determine (Karch et al. 1985; Maeda 2006). Thus, *iab5, iab6, iab7* and *iab8/9* are responsible for the development of the four abdominal segments, A5, A6, A7 and A8/9.

In addition to the spatially collinear arrangement of the modules, the numbers of the modules also corroborate with the number of the segments that they provide identity; the four CRMs drive the identity of four abdominal segments.

Each of the *Abd-B* modules is separated by chromatin domain boundaries in the order *MCP* that separates *iab4* and *iab5* (Mihaly et al. 1998), *Fab6* separating *iab5* and *iab6* (Pérez-Lluch et al. 2008), *Fab7* separating *iab6* and *iab7* (Gyurkovics et al. 1990a), and, *Fab8* that separates *iab7* and *iab8* (Barges et al. 2000). A deletion of several regions of the CRM causes a loss of function of the associated gene that leads to anteriorization of the respective segment (Celniker et al. 1990). For example, the deletion of *iab6* elements leads to a transformation of A6 to A5 (Galloni et al. 1993). The module that becomes activated in a segment, remains activated even in the posterior segment (Bowman et al. 2014; Maeda and Karch 2015). Thus, when a posterior module is not available in case of a deletion, the adjacent anterior module defines the segment identity. Such flies have two copies of A5 followed by A7 (Mihaly et al. 2006). On the contrary, the deletion of boundaries, such as *Fab7*, would lead to the posteriorization of A6 to A7. This is due to ectopic derepression of posterior module *iab7* that prevails over the anterior module *iab6*, and drives the identity of A6. The fly, thus, has A5 followed by two copies of A7 (Gyurkovics et al. 1990b; Mihaly et al. 1998; Iampietro et al. 2010).

Although there are several inversions and a translocation reported for different regions of the BX-C (Celniker and Lewis 1993; Ho et al. 2009; Hendrickson and Sakonju 1995; Stuttgart and Herberth 1988; Gligorov et al. 2018), there are no known cases of targeted duplications or inversions of CRMs concerning their effect on adult segments. Further, many questions concerning the regulation of the BX-C remain unanswered (Bender 2020). For instance, how well do the CRMs rely upon each other for the accessibility of the chromatin to the transcription factors? Is the relative positioning and order of the CRMs important to respond to the upstream spatial cues? What is the relevance of the defined number of modules in the system and can the system accommodate additional modules? The sequential opening of the chromatin represented by the loss of repressive H3K4me3 marks is observed across the BX-C CRMs through anterior to posterior parasegments. If the order or number of CRMs are altered, would the sequential opening model maintain its status-quo?

In order to answer such questions, we generated transgenic lines with FRT insertions in targeted regions across the *Abd-B* CRMs. these lines could be used to create multiple deletions, duplications, and inversions for the desired set of CRMs (Fig. 1A). We modified the *cis*-regulatory landscape of the posterior-most BX-C gene, *Abd-B*, to generate useful resources that will help us advance our understanding of the role of *cis*-regulatory modules in regulating Hox. Further, our study also features a set of *in-vivo* systems to comprehend the arrangement and number of *cis*-regulatory modules in the BX-C. This system can be repurposed to shuffle the arrangement and number of regulatory domains in order to understand the functioning of CRMs in addition to our existing knowledge about the individual regulatory elements.

**Figure 1.**
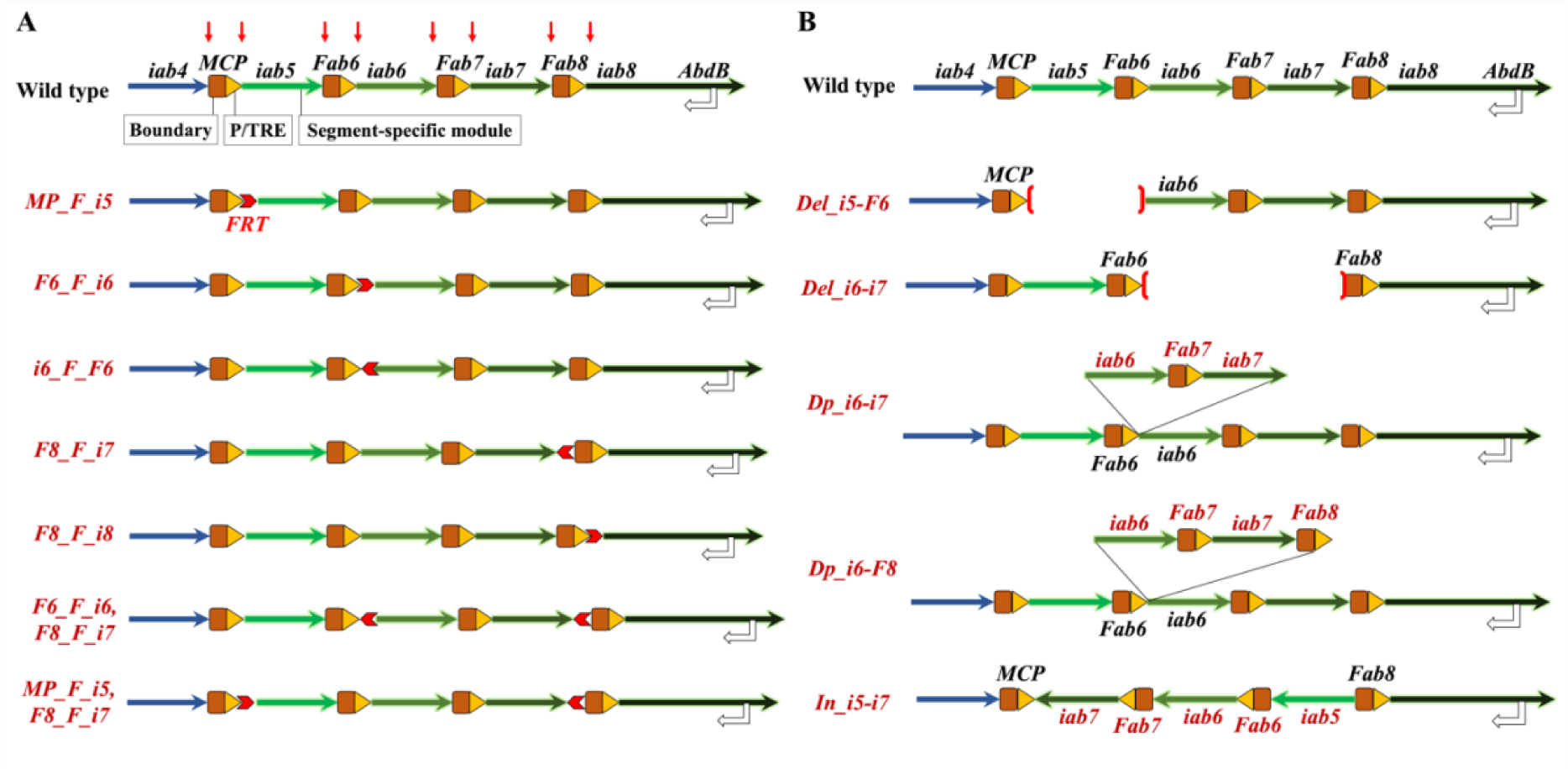
Targeting the cis-regulatory modules of the bithorax complex regulating *Abd-B*: (A) The proposed experiment for the current study is to generate modular deletion, duplication and inversion within the *cis*-regulatory landscape of *Abd-B*. Top row shows the normal arrangement of the CRMs (green solid arrows) of *Abd-B*, iab5, iab6, iab7 and iab8 demarcated by the boundaries MCP, Fab6, Fab7 and Fab8 (brown rectangle and pentagon arrows). The blue arrow depicts iab4 which is a regulator of *abd-A*. The re-arrangement of the CRMs of different nature are shown below the wild-type representation. The concomitant expression of Abd-B driven by the four CRMs in the four abdominal segments of the fly is shown on right. (B) Top row: Representation of the wild-type locus of *Abd-B*. Green arrows are regulators of Abd-B, blue arrow is regulator of abd-A, brown rectangles represent the boundaries and yellow triangles represent the PREs. Red arrows pointing down indicates the sites proposed for FRT insertion for which constructs were generated. Bottom rows indicate the nature of transgenes generated with FRTs (red chevron arrow) in different locations and orientations. (C) The lines with altered CRMs generated in this study.

## Results

### Generation of FRT knock-in transgenes

The BX-C is heavily decorated with binding sites for transcription factors and chromatin remodelers (Négre et al. 2011). Further, a single nucleotide transition of a repressor binding site was shown to have an effect on the ectopic derepression of *iab5* in A3 leading to a prominent A3 to A5 homeotic transformation of the segment (Ho et al. 2009). Thus, the BX-C CRMs need to be tightly regulated and leave narrow margins to make amendments in the genome without affecting *cis*-motifs. Therefore, to insert FRT sites without any further perturbation, we utilized the available data for experimental validations of the BX-C and ChIP data on modENCODE to connote the *Abd-B cis-*regulatory landscape into distinct *cis*-regulatory modules, *iab5* through *iab8* (Fig 1A). The boundaries separating these domains were also annotated based on experimental results from previous studies (Iampietro et al. 2008; Mishra and Karch 1999; Postika et al. 2018; Kyrchanova et al. 2015). Further, regions marking P/TREs distal to the boundaries were annotated based on earlier experimental validations and previously reported PRE mapping tool, PRE mapper (Pérez-Lluch et al. 2008; Mishra et al. 2001; Singh and Mishra 2015; Postika et al. 2021a; Srinivasan and Mishra 2020). The target sites for FRT insertions were selected such that the FRT lands between a boundary and a CRM without perturbing the boundary-PRE combination (Fig. 1A, Supplemental Table S1).

The FRT sequences were cloned in different orientations with required homology arms (Supplemental Table S2) for CRISPR-Cas9 mediated homology-directed repair (Port et al. 2015). The FRTs are inserted between *MCP-iab5*, *Fab6-iab6*, *iab7-Fab8,* and *Fab8-iab8* in different orientations as shown in Fig 1A. For example, the line *MP_F_i5* has FRT inserted between *MCP* and *iab5* with the direction of FRT towards *iab5*. Similarly, the *i6_F_F6* line has an insertion of FRT between *Fab6* and *iab6* with the direction of FRT towards *Fab6*.

The FRT insertion is validated by amplifying the region using primers eccentric to the FRT site. Further, the FRT contains an *Xba-I* restriction digestion site. Thus, the amplified products for positive transgenes are digested with the endonuclease and render two distinct bands on agarose gel electrophoresis (Supplemental Fig. S1, Supplemental Table S6). The FRT insertions are further confirmed by sequencing.

Other than the flies that have FRTs inserted in a single locus, we also generated lines that have a couple of FRTs inserted in the same as well as opposite orientations. The single FRT lines can be used as landing platforms for various other transgenic assays and can also be repurposed to generate targeted deletions and duplications of the CRM-boundary combinations in the locus (Horn and Handler 2005; Phan et al. 2017; Golic and Lindquist 1989). The double FRT with the FRTs in the same orientation can be used to generate deletion in *-cis* or duplication in *-trans*(Golic et al. 1997). The duplicated line can further be used to generate subsequent modules of the *cis*-regulatory landscape by repurposing along with single or double FRTs. On the other hand, the double-FRT line with the two FRTs in opposite orientation can be used to generate inversion of the locus to re-arrange the order of the CRMs and boundary (Golic et al. 1997). Some of the combinations using single FRTs and possibilities of using double FRTs are mentioned in the following sections. A summary of all the FRT lines and genomic rearrangements generated in the study is presented in Fig. 1.

### Deletion of *iab5-Fab6* reveals the non-autonomous function of CRMs

The number of *Abd-B* CRMs corroborates the number of abdominal segments. Thus, we rationalized that the deletion of information about the development of one of the segments can cause an impact on the number of segments formed. Although larger deletions in the region were shown to affect only the identity of segments but not the number, the exact effect in adults is not clear. This is especially because many of the larger deletions including *abd-A* and *Abd-B* CDS lead to embryonic lethality in homozygous conditions. Thus, removing a combination of CRM along with boundary will leave the locus with altered modularity characterized by lesser numbers of CRM-boundary combinations. Towards this, we generated a line that lacks a ∼15.14 kb region spanning *iab5* and *Fab6* (R6.0 – 3R: 16,872,062..16,887,207). Thus, the fly is left with three CRMs instead of four in wild-type animals (Fig. 2A).

**Figure 2.**
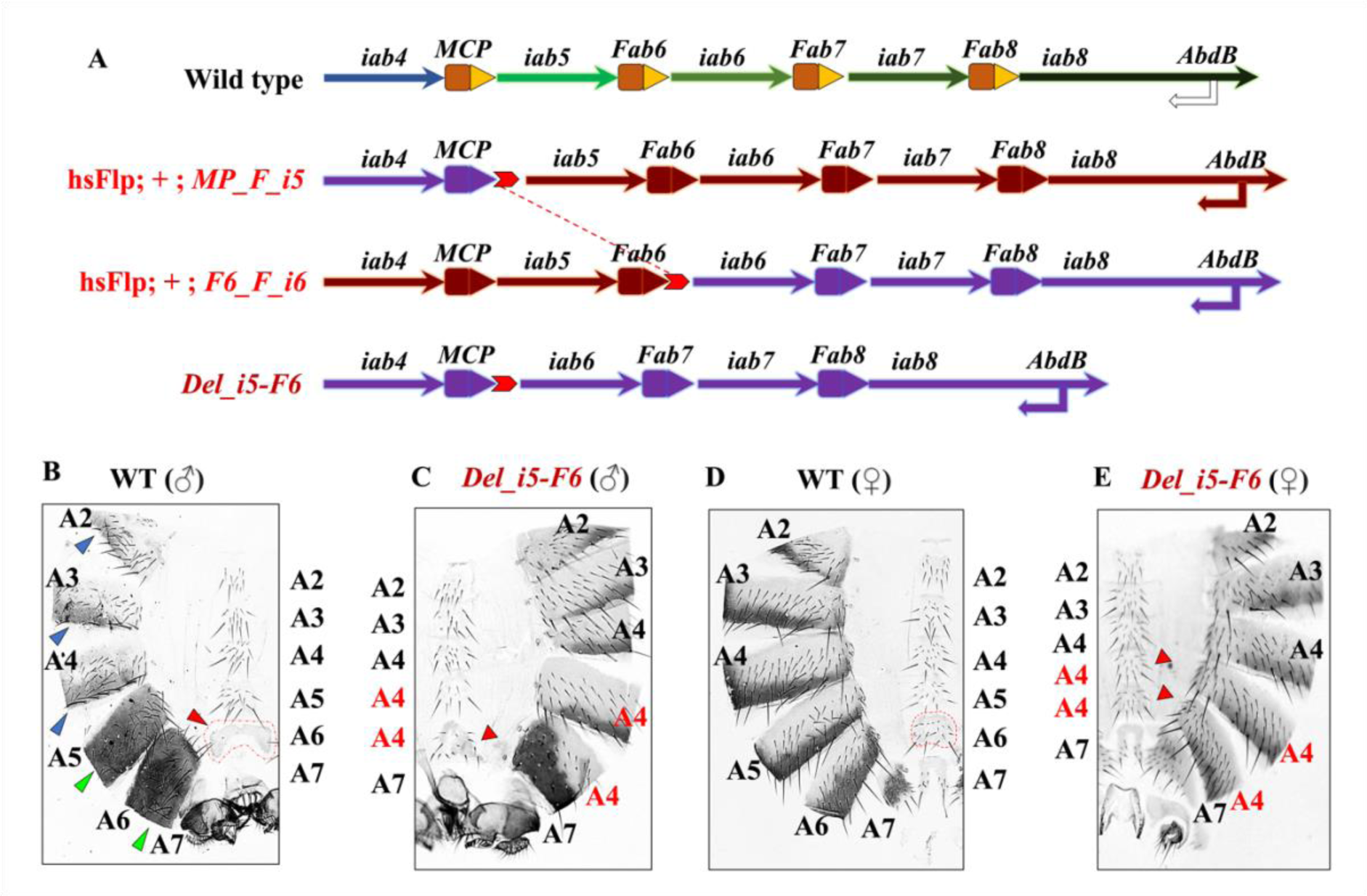
Deletion of *iab5* and *Fab6*: (A) Representation of the wild-type Abd-B cis-regulatory landscape and dichromatic representation of flies with endogenous FRT (red chevron arrow) between *MCP*-*iab5* (*MP_F_i5*), and, *Fab6-iab6* (*F6_F_i6*). The bottom row indicates the deletion product formed as a result of recombination indicated by red-dotted line across the middle rows. (B) Abdominal cuticle preparation of adult male flies two days post eclosion. The curved segments on the left form the dorsal side of the abdomen, while the straight lane of bristles towards the right is the ventral sternites. Each dorsal segment has a distinct pattern of melanisation, as indicated by blue and green arrows. The blue arrows indicate melanisation in the posterior ends of segments A2-A4 under the influence of *abd-A* driven by *iab2* through *iab4*. The green arrows indicate complete melanisation of A5 and A6 influenced by the levels of *Abd-B* regulated by *iab5* and *iab6* in respective segments. Each ventral segment also has a distinct arrangement of bristles. The A6 does not have any bristles on sternite in the wild-type fly, as shown by the red arrowhead and dotted shape. (C) The abdominal cuticle of adult males with deletion of *iab*5 and *Fab6*. The males show a strong homeotic transformation of A5 to A4 characterized by loss of dorsal melanization (curved segments on the right). The flies also show A6 to A4 transformation characterized by loss of melanization of dorsal segment and appearance of bristles on ventral sternites (bold red arrow). (D) Abdominal cuticle preparation adult female flies. The dorsal melanization and the pattern of tergites and sternites are largely indistinct in females. However, the A6 of the females have a hardened sclera in wild-type females (dotted shape). Note that A7 of the wild-type flies have fewer bristles that droop towards the genitalia. The A7 of the fly is also observably smaller than the other segments of the fly. (E) Abdominal cuticle prep of adult *Del_i5-F6* female fly. Note that the sternites in A5 and A6 are strikingly similar to A4 as indicated by bold arrowheads. The tergites and melanisation pattern also appears strikingly similar to A4 in A5 and A6. Thus, both males and females had similar cuticular homeotic transformations in the flies lacking known *iab5* and *Fab6* regions.

To obtain the line lacking *iab5* and *Fab6*, we crossed *hsFlp*; *MP_F_i5* (FRT inserted between *MCP* and *iab5* with the direction of *FRT* towards *iab5*) with *hsFlp*; *F6_F_i6* (FRT inserted between *Fab6* and *iab6* with the direction of FRT towards *iab6*), see Fig 2A. Both lines express *Flippase recombinase* (FLP) under the influence of heat-shock inducible promoter (*hs*). The trans-heterozygous progenies with one allele with an FRT between *MCP* and *iab5* while the other with an FRT between *Fab6* and *iab6* were given heat shock at 37°C for 90 min at an interval of 24 hrs. Heat shocks were given from the late second instar larval stages until the late pupal stages of *Drosophila* development. FLP activity was confirmed as described previously (Pignoni et al. 1997); see methods and supplementary information (Supplemental Fig. S2) for details.

Once the heat-shocked animals eclosed (G1), they were mated with third chromosome balancers to prevent genetic recombination of target loci. The progenies from these crosses (G2) were pooled in batches of 10 and were again crossed with third chromosome balancers. Once enough activity was observed in the vial, the flies were screened via PCR for the desired deletion. The PCR products of flies showing amplification of deletion specific regions were confirmed via Sanger sequencing (Supplemental Fig. S3).

We observed a striking anteriorization of A5 and A6 into copies of A4 in the deletion mutants *Del_i5-F6*. The A5 and A6 in wild-type males are characterized by complete melanization whereas, the A4 has melanization in the posterior end. The homozygous fly for the deletion of *iab5-Fab6* shows a transformation of A5 to A4 characterized by the loss of melanization. Moreover, even the A6 was transformed into a copy of A4 characterized by the loss of melanization and appearance of ventral bristles which are otherwise absent in a wild-type male (Fig. 2B-C).

Previous studies showed the functioning of CRMs in an autonomous manner wherein the autonomy is largely maintained by the boundary elements demarcating the modules (Hagstrom et al. 1996; Maeda and Karch 2015). Deletions in *iab6* or *iab7* do not affect the functioning of each other in the associated segments A6 or A7. For instance, the deletion of *iab6* regions would cause A6 to transform into a copy of A5 but does not have an impact on A7 or A8 (Iampietro et al. 2010). On the other hand, the deletion of a boundary would ectopically activate the posterior CRM. For example, the deletion of *Fab6* would lead to the transformation of A5 into a copy of A6. Akin to that, the deletion of *Fab7* will transform A6 into A7 (Mihaly et al. 1998). Therefore, we predicted that the deletion of *iab5-Fab6* will lead to the transformation of A5 into a copy of A6, a phenotype largely dominated by the deletion of *Fab6*.

In contrast to our expectations, we observed both A5 and A6 transforming into copies of A4 in males and females (Fig 2B-E). Since the deletion of *Fab6* does not impact *iab6* activity, we reasoned that the regions in *iab5* are responsible for *iab6* activation, thereby defying the autonomy of *iab6*. Interestingly, Postika et al. reported a similar effect observed due to deletions of several regions within *iab5*. Since the deletion we generated encompassed all the regions mentioned by Postika et al., we see a similar phenotype with a clean transformation of A5 and A6 into copies of A4 (Postika et al. 2021b, 2021a). Thus, although the system is left with modules responsible for the formation of A6 through A8; the activators in anterior CRM, *iab5*, are required for the execution of the spatial code in A6 but not in A7 or A8.

### Deletion of *iab6-iab7* transforms A6 and A7 into copies of A5

The transgenic assays and genetic interactions from previous studies have revealed that boundaries interact with each other (Maeda and Karch 2007; Kyrchanova et al. 2011; Singh and Mishra 2015). Therefore, we deleted a region spanning *iab6* through *iab7* thereby juxta positioning *Fab6* next to *Fab8* (Fig. 3A). Towards this, we generated a line with two FRTs inserted in the same orientation in two different loci within *Abd-B cis*-regulatory landscape. One of the FRTs was inserted between *Fab6* and *iab6* with the direction of FRT towards *Fab6* (*i6_F_F6*). The other FRT was knocked-in between *iab7* and *Fab8* with the orientation of FRT towards *iab7* (*F8_F_i7*) as shown in Fig. 3A. Although the FRTs were inserted in *cis*- and a homozygous fly could have rendered us the required deletion, we are also open to the possibility of the recombination to have occurred in *-trans* as indicated in Fig. 3A.

**Figure 3.**
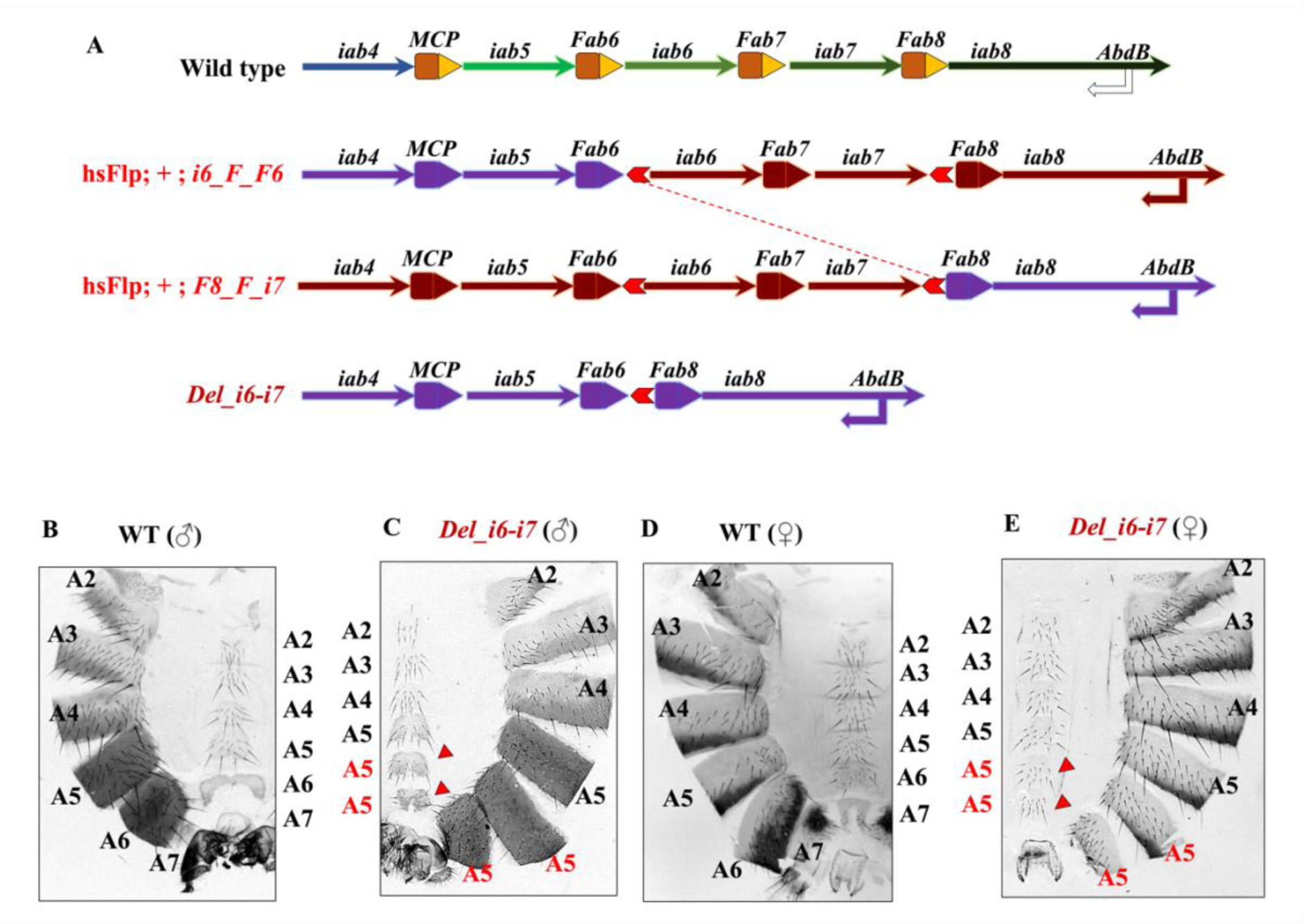
Deletion of *iab6-iab7*: (A) Representation of the wild-type *Abd-B* cis-regulatory landscape and dichromatic representation of flies with endogenous FRT (red chevron arrow) between *Fab6*-*iab6* (*i6_F_F6*), and, *iab7-Fab8* (*F8_F_i7*). The bottom row indicates the deletion product formed as a result of recombination indicated by red-dotted line across the middle rows. (B-E) Abdominal cuticle preparation of adult flies two days post eclosion. (B) Abdominal cuticle preparation of wild-type male. (C) The abdominal cuticle of adult males with deletion of *iab*6 through *iab7*. The flies show a strong homeotic transformation of A6 and A7 to copies of A5. The A6 to A5 transformation is characterised by broadened A6 and sternal bristles’ appearance (bold red arrow). The A7 to A5 transformation is characterised by a distinct segment with dorsal melanisation similar to A5. The transformed segment also has sternal bristles similar to A5. (D) Abdominal cuticle preparation of wild type female. (E) Abdominal cuticle prep of adult *Del_i6-i7* female fly. Note that the sternites in A6 and A7 are strikingly similar to A5 (bold red arrows). The tergites and melanisation pattern also appears strikingly similar to A5 in A6 and A7. Thus, both males and females had similar cuticular homeotic transformations in the flies lacking known *iab6* through *iab7* regions.

The FLP mediated recombination under the influence of a heat-shock inducible promoter was carried out as described in the previous section. To molecularly validate the deletion, the genomic DNA of the putative mutants was isolated and amplified using a combination of primers specific for the deletion locus upon recombination; an expected juxta positioning of *Fab6* and *Fab8*. The repositioning would amplify a specific product with the forward primer for screening FRT between *Fab6* and *iab6* (F6i6_ScrF) and the reverse primer for screening FRT between *iab7* and *Fab8* (i7F8_ScrR). A resultant product of 974 bp confirms the deletion event as shown in Supplemental Fig. S4. The endogenous locus at the approximate junction of *Fab6* and *iab6* was chosen as a control locus. This region was amplified in the genomic DNA of a wild-type CS fly but not the deletion mutant (Supplemental Fig. S4C). The amplified product for deletion mutants was confirmed by sequencing (Supplemental Fig. S4D). The total length of the deleted region spans 31.374 kb from 3R: 16,887,208 to 3R: 16,918,581 (Genome assembly R6.0).

We observed a phenotype as was expected with the deletion of the CRMs *iab6* and *iab7* that are responsible for determining the identities of A6 and A7. There is a complete loss of function in the homozygous flies carrying the desired deletion as characterized by the complete transformation of A6 and A7 into copies of A5 (Fig. 3B-E). Additionally, the males also show genital rotation of varying degrees and are sterile. The deletion of intermittent boundary *Fab7*, and, juxtapositioning of *Fab6* and *Fa8* do not cause an effect on the functioning of *iab5*. The phenotype is also apparent in females wherein a wild-type female has 8-10 narrow groups of drooping bristles in A7. In the mutant females, A7 is broader, and a complete set of sternites appear on the ventral surface of the segment (Fig. 3D-E). The juxta positioning of *Fab6* and *Fab8* does not seem to have an effect on regulation of *Abd-B* via *iab5*.

### Duplication of *iab6-iab7* renders phenotype dominated by the posterior module *iab7*

The number of CRMs in the BX-C is equivalent to the number of segments that they provide identity. Hence, it is important to understand the functional correlation of the number of CRMs and their role in providing identity to respective segments. Towards this, we generated a mutant line with duplication in *iab6* through *iab7* such that the modularity is lost. The duplication spans 31.374 kb (3R: 16,887,208..16,918,581). This arrangement juxtaposes an additional copy of *iab6* distal to *iab7* (Fig. 4A) without a boundary. Using the double FRT line used for obtaining the deletion of *iab6-iab7*, we generated a recombinant line with the duplication of the same region as indicated in Fig. 4A.

**Figure 4.**
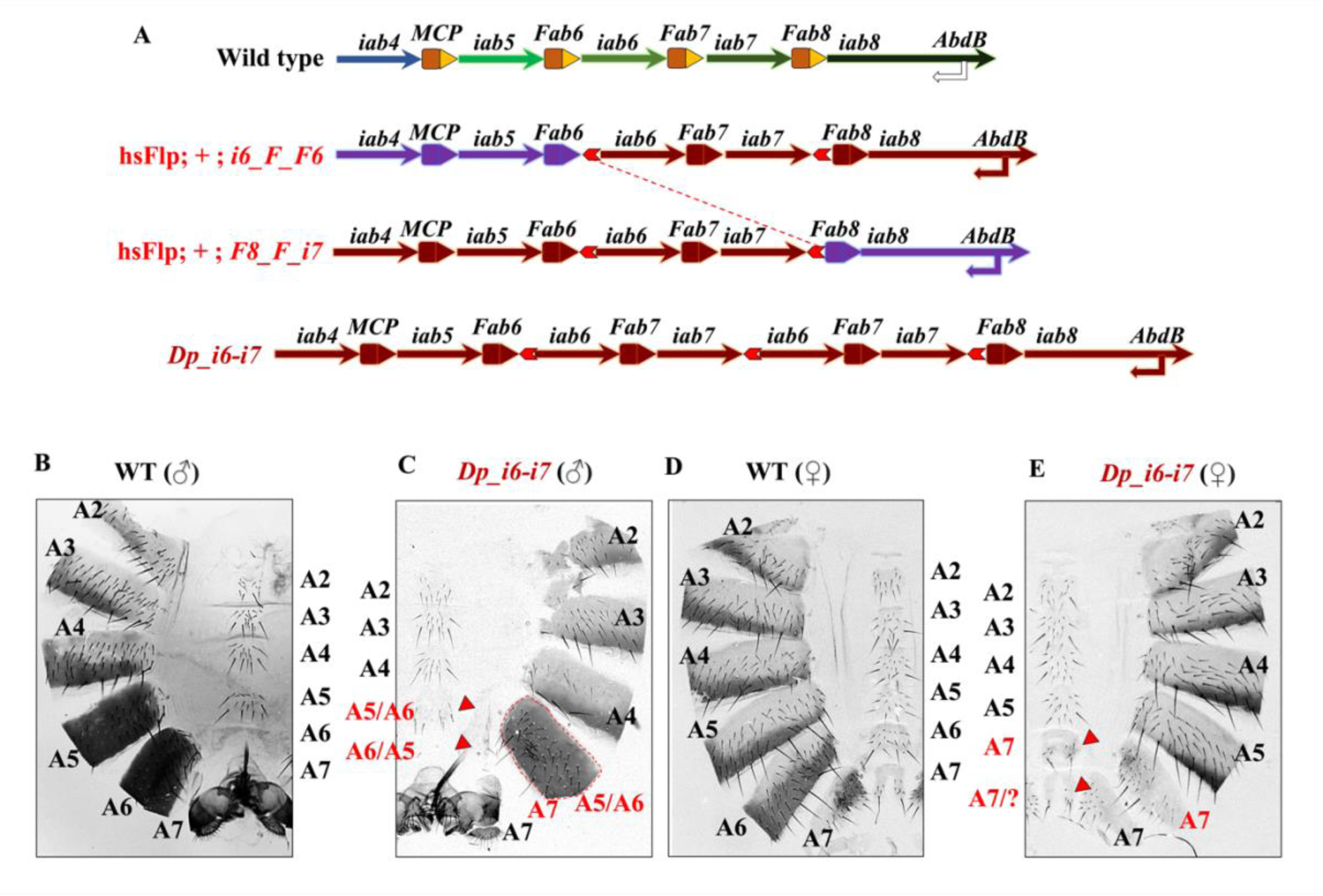
Duplication of *iab6* through *iab7*: (A) Representation of the wild-type *Abd-B* cis-regulatory landscape and dichromatic representation of flies with endogenous FRT (red chevron arrow) between *Fab6*-*iab6* (*i6_F_F6*), and, *iab7-Fab8* (*F8_F_i7*). The bottom row indicates the duplication product formed as a result of recombination indicated by red-dotted line across the middle rows. (B-E) Abdominal cuticle preparation of adult male flies two days post eclosion. (B) Abdominal cuticle preparation of wild-type male. (C) The abdominal cuticle of adult males with duplication of *iab*6 through *iab7*. The flies show a strong homeotic transformation of A6 to a copy of A7. The transformation is characterised by loss of segment with dorsal melanization. The A6 also has partial signatures of anteriorization indication by the appearance of sternal bristles and loss of sclera. (bold red arrow). The A5 of the fly also shows partial posteriorization indicated by loss of sternal bristles. The partial posteriorization is also suggested by slight narrowing of dorsal abdomen akin to A6 of a wild-type fly (shown by the red dotted peripheral structure). (D) Abdominal cuticle preparation of adult female flies. (E) Abdominal cuticle prep of adult *Dp_i6-i7* female fly. The A5 of the fly appears normal. The A6 shows a clear homeotic transformation into a copy of A7 by the appearance of A7-specific sternal bristles. Note that the sternites in A6 and A7 are strikingly similar (bold red arrows). The tergites and melanisation patterns of A6 and A7 also appears strikingly similar. Thus, both males and females had similar cuticular homeotic transformations in the flies lacking known *iab6* through *iab7* regions. The A7 of the flies show loss of melanization, indicating either transformation into A8 or uncharacterised structures.

To obtain the duplication line, we screened the recombinants from the progenies of flies set for obtaining the deletion of *iab6* through *iab7* in the previous section (Supplemental Table S9-S10). The expected recombination of the two loci in *-trans* is depicted in Fig. 4A with dichromatic representations of the parent and prospective recombinant alleles. Note that the duplication allele has three FRTs. One of the FRTs is derived from recombining FRTs between *Fab6-iab6* and *iab7-Fab8*. The other FRTs are present from the parent allele. This line is a novel playground to recombine various transgenic animals having FRTs at different loci in the *Abd-B cis*-regulatory landscape. The three FRTs can be repurposed differently to alter the modularity of the locus in varied manners.

For the molecular validation, the genomic DNA of putative recombinants was isolated and amplified by primers specific to the duplication locus. The forward primer to screen FRT between *iab7* and *Fab8* (i7F8_ScrF) and the reverse primer to screen FRT between *Fab6* and *iab6* (F6i6_ScrR) amplified a specific 534 bp product indicating the presence of duplication allele. The endogenous *Fab6-iab6* and *iab7-Fab8* loci were used as controls. The amplified products were confirmed using sequencing (Supplemental Fig. S5).

The fly with the duplication of *iab6* through *iab7* (*Dp_i6-i7*) has multiple changes in the arrangement of modules. The fly has two *iab6* modules, one flanked by *Fab6* and *Fab7* as in wild-type conditions, and the other is juxtaposed next to *iab7* without intermittent boundary. We observe that there is a partial posteriorization of A5 in adult males as suggested by the decrease in the number of bristles in sternites and partial morphological changes in the dorsal A5 as indicated by dotted shape in Fig. 4C. Further, the A6 identity is completely transformed into a copy of A7 towards the dorsal side. However, on the ventral end, A6 shows a partial anteriorization as suggested by the presence of a few bristles as well as loss of sclera formation. There are no apparent changes in the male A7 of the transgenic animals despite two copies of *iab7*. A similar effect of A6 to A7 transformation is also evident in females (Fig. 4B-E).

The presence of extra modules was particularly interesting to understand the effect of relative positioning of *iab6* and i*ab7*. We thus expected a mixed gain and loss of function of A6 as well as A7. We were also open to the possibility of observing a segment with novel features since the duplicated *iab7-iab6* formed a larger module flanked by *Fab7*. Moreover, the presence of extra modules would also change the relative positioning of *iab5* with respect to the *Abd-B* promoter and thus an effect in A5 was also expected. However, a strong A6 to A7 and further posteriorization of A5 indicate the dual copies of *iab7* were dominant over the two copies of anterior module *iab6* as well as the wild-type *iab5* irrespective of relative positioning.

### Duplication of *iab6-Fab8* alters modularity and renders variable phenotype

In the previous duplication, the modularity was perturbed due to the absence of a boundary element between *iab7* and *iab6*. Therefore, to preserve modularity, we duplicated the 34.776 kb (3R: 16,887,208..16,921,983) region from *iab6* through *Fab8*, keeping the modularity intact. Thus, each CRM is still flanked by a combination of boundary-PRE as shown in Fig. 5. Further, to keep one of the CRMs as an internal control, we left *iab5* intact and duplicated *iab6* through *Fab8*.

**Figure 5.**
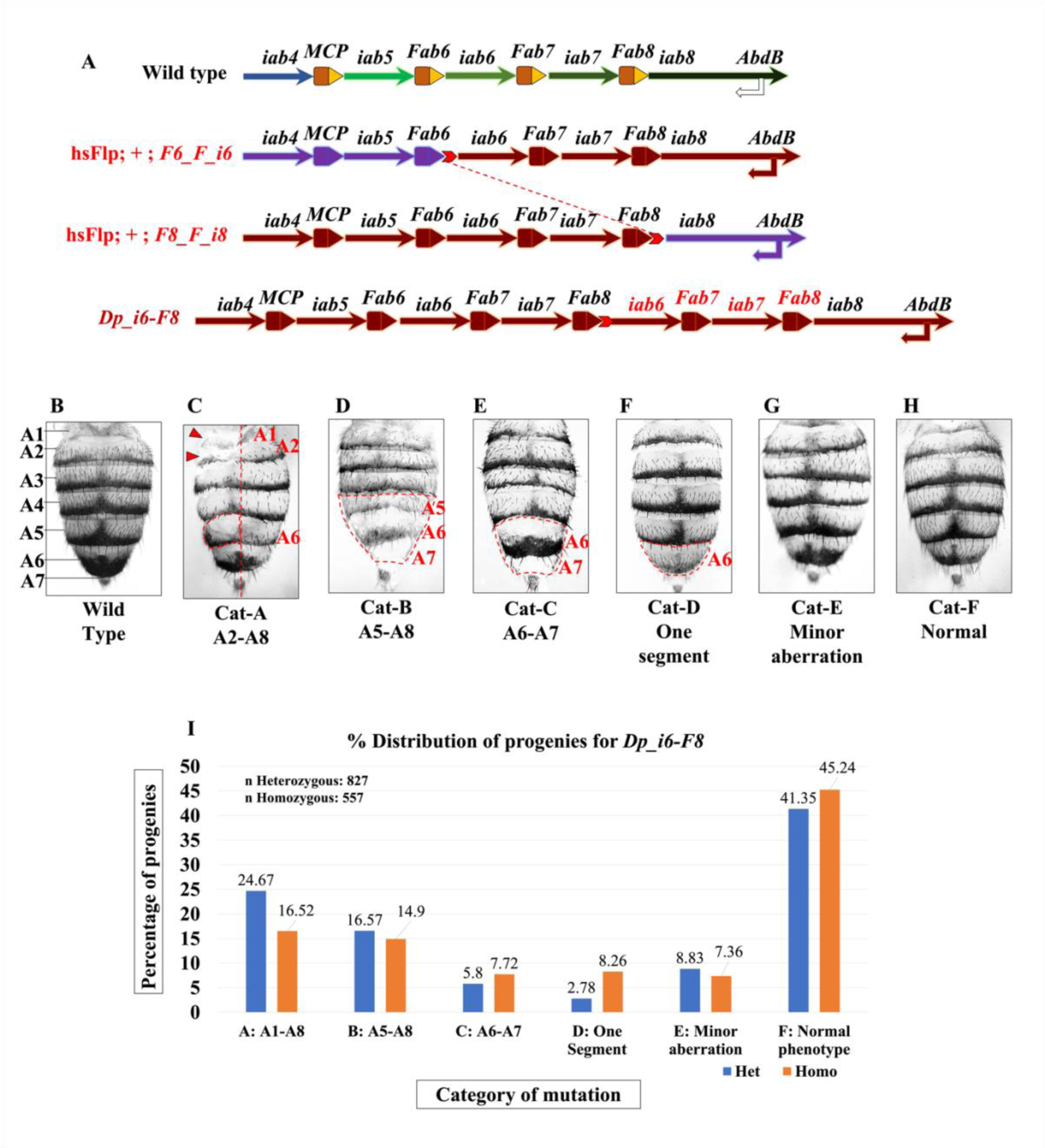
Duplication of *iab6* through *iab7*: (A) Representation of the wild-type *Abd-B cis-*regulatory landscape and dichromatic representation of flies with endogenous FRT (red chevron arrow) between *Fab6*-*iab6* (*i6_F_F6*), and, *Fab8-iab8* (*F8_F_i8*). The bottom row indicates the duplication product formed as a result of recombination indicated by red-dotted line across the middle rows. (B-H) Bright-field images of the dorsal abdomen of two-days post eclosion adult females from different classes of mutants. The images are represented in grayscale for easier visualization of the segmental pattern. (B) A wild-type CS fly is shown as a control for patterning defects in the mutants. (C-H) Flies with variable phenotype classified into different categories depending upon the extent of segmental aberration. See text for details. (I) A standard histogram plotted for the distribution of 827 heterozygous and 557 homozygous flies for the duplication of *iab6* through *iab8* with different phenotypes grouped under categories A-F (see text for details). Blue bars indicate the percentage distribution of heterozygous flies, and the orange bars represent the percentage distribution of the homozygous flies.

To obtain a fly with proposed duplication, we genetically crossed the flies with FRT insertions between *Fab6-iab6* and *Fab8-ia8* to obtain a trans-heterozygous line. These flies were in the background of *Flp* recombinase expressed under the control of a heat-shock inducible promoter as described earlier. A brief schematic of flies used and the duplication upon recombination is depicted in Fig. 5A. For details, see methods and Supplemental Table S9-S10.

Next, to validate the duplication event molecularly, the genomic DNA of prospective recombinants was isolated and amplified by primers specific to the duplication locus (F8i8_ScrF and F6i6_ScrR). The intact loci at *Fab6-iab6* and *Fab8-iab8* were amplified as controls. Only the flies possessing the desired duplication showed an amplified product (Supplemental Fig. S6C). Genomic DNA from CS flies was used as a control. Unlike the duplication locus that was amplified only in the positive recombinants, the *Fab6-iab6* and *Fab8-iab8* loci were amplified in both the duplication and CS genomic DNA. The products were later confirmed by sequencing (Supplemental Fig. S6C-D).

In the *Dp_i6-F8* flies, the combination of CRM-boundary is intact. Thus, we expected a mixed gain and loss of functions in A6 and A7. Since *iab6* and *iab7* are regulators of *Abd-B*, an effect could also have been obtained in A5 and A8. The extra copies of *iab6* and *iab7* were modularly flanked by *Fab6* and *Fab7* akin to the wild-type scenario. Surprisingly, we observed a variety of phenotypes with perturbations ranging from disturbances in A1 to a normal-looking fly similar to wild-type (Fig. 5B-H).

We classified the phenotypes into six different categories as follows: Category A (Cat-A) mutants show the most extreme phenotypes with all abdominal segments from A1 to A8 severely perturbed. Several flies have disturbances in left-right or dorsal-ventral patterning. These flies are often sterile (Fig 5C). Cat-B animals showed disturbances in segments A5 to A8 (Fig. 5D). The rationale for this category is the role of *iab6* and *iab7* to regulate *Abd-B* that defines A5-A8. Any segment perturbed anterior to A5 was not counted in this category, and instead, was noted in the previous category, A. The Cat-C mutants displayed disturbances in A6 as well as A7 (Fig. 5E). Since the duplicated *iab*s are responsible for providing identities to A6 and A7, flies having perturbations in only these segments were considered a separate category. Cat-D mutants had a perturbation in only one of the abdominal segments. Often these perturbations were restricted to A6 or A7 (Fig. 5F). In the next category, Cat-E mutants display a slight perturbation of dorsal melanization and often rendered “wavy” segmental boundaries (Fig. 5G). The Cat-F mutants show a wild-type phenotype with no observable change in the cuticle or ventral bristle patterning (Fig. 5H).

Both males and females showed the mentioned pattern. However, females were better to observe via imaging owing to clear demarcations of seven abdominal segments followed by genitalia. Also, the cuticles of these animals were extremely brittle, and therefore, the abdomen was directly imaged under the stereomicroscope (Fig. 5B-H; see methods for details).

All the classes of mutants were confirmed to have the same genotype molecularly. Further, the inbreeding of mutants from all the phenotypes, including Cat-F (normal segments), rendered progenies with varying phenotypes again. The development of progenies with inconsistent phenotypes is true for heterozygous and homozygous flies from these categories. Hence, we selected only category F adults and set up a breeding experiment to check the distribution of progeny with different phenotypes.

We observed almost half the population for homozygous (45.24%) and heterozygous (41.35%) progenies were normal (Cat-F). In contrast, the remaining categories had a smaller number of flies in the population (Fig. 5I). Interestingly, ∼25% heterozygous (24.67%) and ∼17% homozygous (16.52%) flies showed phenotypes grouped under category A. The heterozygous flies showed 1.5 times a greater number of progenies belonging to Cat-A than homozygous (Fig 5I). Our results suggest that more heterozygous flies tend to be severely affected by the duplication of CRMs (Supplementary Data).

Thus although the duplication of *iab6-Fab7* perturbs the genomic modularity of the locus (previous section), the segmental boundaries are intact. However, duplication of modules keeping genomic modularity intact (*Dp_iab6-Fab8*) disrupts the segmental boundary in a manner that requires further investigations to understand the BX-C better.

### The inversion of *iab5* through *iab7* provides mixed and altered identities of the associated segments, A5, A6 and A7

The CRMs of the BX-C are present in a spatially colinear manner. The three CRMs, *iab5*, *iab6* and *iab7* regulate *Abd-B* to provide identities to A5, A6 and A7 respectively. The specific ordering of the CRMs can be consequential towards co-regulation of the associated gene (Mateo et al. 2019; Maeda and Karch 2015; Hajirnis and Mishra 2021). One way to test this hypothesis is to invert the order of these regions. Towards this, we generated a 46.520 kb (3R:16,872,062..16,918,581) inversion of *iab5* through *iab7* (Fig 6).

**Figure 6.**
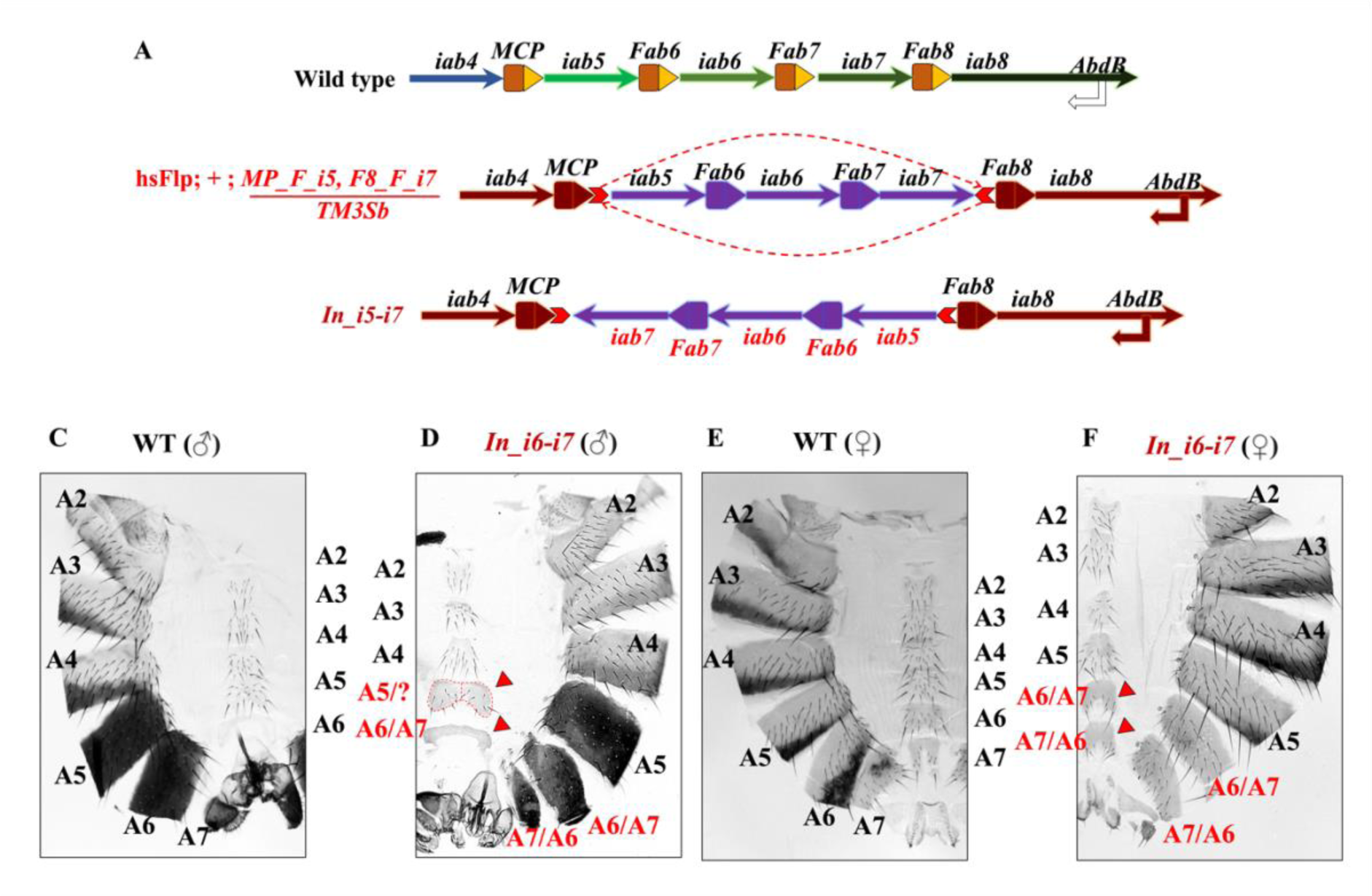
Inversion of *iab5* through *iab7*: (A) Simple representation of flies with endogenous FRT (red chevron arrow) between *MCP* and *iab5* with the direction of FRT towards *iab5* (*MP_F_i5*). The flies also have another FRT inserted between *iab7* and *Fab8* with the direction of FRT towards *iab7* (*F8B_F_i7*). The activity of FLP causes an inversion of iab5 through iab7 as shown in the bottom row. The FRT mediated recombination happens across the two FRTs in *cis-* as indicated by curved dotted arrows in the middle rows. (B) Cuticle preparation of adult male abdomen. (C) The abdominal cuticle of adult males with inversion of *iab5* through *iab7*. The flies show a strong homeotic transformation of A6 to a copy of A7. The transformation is characterised by loss of segment with dorsal melanization. The A6 also has partial signatures of anteriorization indicated by the appearance of sternal bristles and loss of sclera (bold red arrow). The A5 of the fly also shows partial posteriorization indicated by loss of sternal bristles. The partial posteriorization is also suggested by slight narrowing of dorsal abdomen akin to A6 of a wild-type fly (shown by the red dotted peripheral structure). (D) Abdominal cuticle preparation and mounting of 2 days post eclosion adult female flies. (E) Abdominal cuticle prep of adult *Dp_i6-i7* female fly. The A5 of the fly appears normal. The A6 shows a clear homeotic transformation into a copy of A7 by the appearance of A7-specific sternal bristles. Note that the sternites in A6 and A7 are strikingly similar (bold red arrows). The tergites and melanisation patterns of A6 and A7 also appear strikingly similar.

We generated a fly having two FRTs in the opposite orientations inserted between two loci: *MCP-iab5* and *iab7-Fab8*. The FRT inserted between *MCP-iab5* was oriented towards *iab5*, while the FRT inserted between *iab7* and *Fab8* was oriented towards *iab7*. These flies were crossed with a third chromosome balancer, and the heterozygous larvae were given heat shock as described earlier (Supplemental Table S11-S12). The eclosing adults post-heat shock (G0) were crossed with balancers in anticipation that their gametes had undergone the desired recombination (Fig. 6A). The progenies (G1) emerging from these crosses were further crossed with third chromosome balancers. The G1 flies, after three to four days of mating, were sacrificed for molecular screening.

To molecularly screen for the inversion event, the forward primer for *MCP-iab5* locus (Mi5_ScrF) and the forward primer for *iab7-Fab8* locus (i7F8_ScrF) were repurposed. The forward primer for the *iab7-Fab8* locus is now used as a reverse primer for *MCP-iab7* recombined locus. Similarly, the recombined locus of *iab5-Fab8* was screened by repurposing the reverse primer to screen FRT at *MCP-iab5* (Mi5_ScrR) locus with the reverse primer to screen FRT at the *iab7-Fab8* region (i7F8_ScrR) as explained in Supplemental Fig. S7. A specific product upon repurposing the primers is obtained only for the flies that had an inversion event. The endogenous loci of *MCP-iab5* and *iab7-Fab8* were amplified only in the CS fly and not in the inversion line. The recombined loci were further confirmed by sequencing (Supplemental Fig. S7).

An expected outcome of this inversion was a wild-type phenotype for the mutant animal if all the elements are perfectly exchangeable. The other direct possibility was that the “code” for segmental identity resides in the order of CRMs, and therefore reversing their order might reverse the identities of abdominal segments. In this case, we expect a fly that forms A4, followed by A7, A6 and A5 in the anteroposterior order. However, we noted flies with different phenotypes for different segments (Fig 6B-E).

We must also consider that the order of CRMs was reversed, and their relative positioning was also exchanged. For instance, the native position of *iab5* now has the presence of *iab7* while *iab5* replaces *iab7* as it is juxtaposed next to *Fab8* (Fig. 6A). On the contrary, the *iab6* is in the same position as before but is present in an opposite orientation. The re-orientation of *iab6* also includes the change in directionality of flanking boundaries *Fab6* and *Fab7*. Notedly the boundaries function in an orientation-dependent manner (Zhou et al. 1996; Hogga and Karch 2002; Postika et al. 2018). Therefore, altering the relative positioning of boundaries can also contribute to the phenotypes obtained.

The A5 segment of the flies shows partial gain and loss of function. The numbers and density of ventral bristles decrease, indicating a gain of function to A6 (Fig 6C-D). Occasionally, several males also show partial loss of melanization in A5, indicating partial transformation into A4. The mixed gain and loss of function phenotypes indicate a cell-type-specific behavior of the modules within a particular segment of the mutant animal. The ventral A5 also develops a sclera similar to A6, indicating slight posteriorization (Fig. 6D). The PREs associated with the boundaries in the BX-C provide temporal and cell-type-specific regulation of the adjacent CRMs (Ringrose and Paro 2004). The absence of any major PRE associated with a boundary near *iab5* partly explains the incomplete anterior- and posteriorization upon inversion.

Next, even though *iab6* is in the same position relative to wild-type condition, it shows a partial gain of function transformation into A7 as indicated by narrowing of the dorsal melanization and sternite in the males (Fig. 6C). The effect is also clearly observed in A6 of the females wherein the ventral bristles attain the identity of the A7 (Fig. 6D-E). Notedly, although the *iab6* remains in the same relative order concerning adjoining boundaries and CRMs, the direction of the module is opposite. It has been previously shown that the direction of the BX-C boundaries including *Fab6* and *Fab7* is crucial to performing the insulator bypass activity (Zhou et al. 1996; Postika et al. 2018). In the inversion mutant, since both the boundaries are inverted, *iab6* is misregulated. The studies on orientation were done individually on different elements. The rearrangement in the inversion locus is a complex interplay between many of them. Therefore, the mutants should be probed deeper, especially for the chromatin conformations of the CRMs to understand the interaction of different elements upon inversion.

Further, the A7 shows an opposite transformation when compared to A6. A distinct dorsal melanized segment is present in males in the position of A7, thus indicating anteriorization of the segment into a copy of A6 (Fig. 6C). However, the segment is not completely transformed into A6. A similar effect is evident in the A7 of adult females by the appearance of extra bristles and broadening of the sclera (Fig. 6E). The male flies also have a genital rotation of varying degrees and is not consistent. Additionally, homozygous flies with the inversion of *iab5* through *iab7* are sterile.

As mentioned earlier, an inversion such as this includes multiple disruptions of the regulatory landscape of the associated gene and although individual elements have been dissected earlier, the complex interplay of multiple elements is interesting to probe in such scenarios.

## Discussion

Homeotic genes or *Hox* code for sequence-specific transcription factors that define the anterior-posterior body axis of a developing bilaterian embryo (Hajirnis and Mishra 2021). The *Hox* genes were discovered in *Drosophila melanogaster* wherein they display a striking property of spatial collinearity (Lewis 1978). That is, the genes are arranged in a complex and, the genes that are present at one end of the complex are responsible for the development of the anterior part of an embryo while the genes present at the other end are responsible for the development of the posterior regions. The posterior *Hox* complex in *Drosophila melanogaster*, the bithorax complex presents us with a unique opportunity to study the relevance of the higher-order arrangement of the *cis*-regulatory modules (CRMs) of the homeotic genes (Maeda 2006). The three genes of the BX-C: *Ubx*, *abd-A* and *Abd-B* are regulated by nine *cis-*regulatory modules *abx/bx* and bxd/pbx regulating *Ubx* in posterior T2 through A1, *iab2* through *iab4* regulating *abd-A* in A2 through A4, and, *iab5* through *iab8* regulating *Abd-B* in A5 through A8 (Maeda and Karch 2015). The *iab*s are arranged in a spatially collinear manner with respect to the segments that are affected upon mutating them. Also, the numbers of the *iab*s corroborate with the number of abdominal segments formed in the fly. In the present study, we generated a set of FRT transgenes to alter the *cis*-regulatory landscape of the posterior-most *Hox* gene *Abd-B*. These lines can be used independently to insert test elements in the fly or can be repurposed to alter the regulatory landscape in *Drosophila*.

The role of many of the *cis*-regulatory elements or modules is known in their native structure (Iampietro et al. 2010, 2008; Zhou and Levine 1999; Cléard et al. 2006). Notedly, many of the existing studies regarding genome manipulation of a similar kind are limited by the status-quo of the targeted manipulations, that is, the alterations lack the flexibility to incorporate novel features or re-shuffle the existing modules (Li et al. 2015; Guo et al. 2015; Fabre et al. 2017). In the current study, the use of FLP-FRT at modular junctions makes the system more dynamic and robust. The alterations in the genome are quickly interchangeable. In fact, in the case of duplication of *iab6-iab7*, the fly has three FRTs and thus it can be repurposed to develop a plethora of downstream combinations of modules. For instance, if the *iab7-iab6* fused domain is behaving like a “super-module”, an additional such module can be introduced in the system using FLP mediated recombination using the lines we generated. Furthermore, each of the lines can be followed individually in different directions. For example, it is interesting to probe the chromatin landscape and chromatin interaction in the fly bearing an inversion of *iab5* through *iab7*. Previous studies have shown sequential derepression of the BX-C *cis-*regulatory modules from anterior to posterior segments; a feature called open for business model of the BX-C regulation (Bowman et al. 2014; Maeda and Karch 2015). However, one key aspect to probe in the sequential opening model is the ability of these modules to sense the spatial cue of accessibility (Kyrchanova et al. 2015). Therefore, in the case of inversion, segment-specific methylation marks would reveal novel aspects of sequential opening. Moreover, understanding segment-specific interactions of enhancers and promoters via techniques like ORCA and Hi-C will shed light upon the functioning of these modules in the altered scenario (Mateo et al. 2019).

In cases such as *Dp_iab6-Fab8*, where the modularity is intact with respect to the positioning of CRMs and boundaries but the number of modules has been altered; the variability in phenotype is interesting to probe. For instance, the variability can arise from the change in dosages of the non-coding RNAs (ncRNA) generated from the *cis*-regulatory locus of *Abd-B* (Garaulet and Lai 2015; Gummalla et al. 2014). Or, the variability could be caused by ectopic transvection happening in different cell types in a stochastic manner (Vazquez et al. 2006; Hendrickson and Sakonju 1995). One way to probe the latter is to investigate the role of epigenetic modifiers such as PcG and trxG proteins to modulate the phenotypic variability (Singh and Mishra 2015). Genes like *Zeste* are known players of transvection (Sipos et al. 1998; Birve et al. 2001). The effects of mutations in such genes can be probed to understand the nature of variability present in the system. The other possibility pertaining to the differential dosages of ncRNAs can be probed by *in-situ* hybridization (Arib et al. 2015). However, the definitive changes in the ncRNAs corresponding with the adult phenotypes require further investigation. With the advancement of genomics and gene-editing techniques, many such mutations as presented in the current study can be generated to understand the evolutionary significance of the CRMs and *Hox* arrangement in different organisms. Additionally, comprehending the epigenomic landscape and DNA looping becomes important to grasp the depths of CRM functioning in such conditions.

In summary, we generated a set of transgenic flies that can alter the modularity of *cis-*regulatory domains in the BX-C of *Drosophila melanogaster* and open avenues to explore the fundamental basis of body axis formation. The novel and unanticipated phenotypes obtained in the study clearly demonstrate that our current understanding of the mechanisms of regulation of bithorax complex is substantially inadequate. This study also shows the potential of targeted re-arrangement of the modules to elucidate the role of the genomic landscape of the bithorax complex. These resources will shed light on important aspects of *Hox* regulation including but not limited to finer details of the order of the CRMs, chromatin conformations, enhancer-promoter interactions, *modus operandi* of regulatory boundaries, or causal relation between histone modifications and expression of associated Hox. Genetic interactions of the novel modules will provide crucial insights underlying their concerted functioning and open a new paradigm for the business of the bithorax complex.

## Materials and Methods

### Primer designing for FRT insertion

The primers for targeting Cas9 were obtained after submitting query sequences for different regions of the BX-C at http://crispor.tefor.net/. *Drosophila melanogaster* BDGP release 6 (R6.0) was selected as the reference genome, and 20bp crRNA with *S. pyogenes* Cas9 5’-NGG3’ PAM was curated for obtaining targets. From the list of suitable gRNAs, sequences with an MIT specificity score of 97 or more (preferably 100), CFD specificity score of 97 or more (preferably 100) and a Lindel score of 80 or more were selected as preferred guides. Due care was taken to select primers with the least number of off-targets as indicated in the relevant column.

Upon successful selection of guides, 1000-1200 bp upstream and downstream region of the cut site was selected as a query to design primers for amplification and cloning of homology arm for donor constructs in https://www.ncbi.nlm.nih.gov/tools/primer-blast/. For all the FRT insertions, the centromeric proximal homology arm to the cut site was denoted as the left homology arm (LH), while the distal homology arm was denoted as the right homology arm (RH). The reverse primer for LH was selected as 20-24 bp immediately centromeric proximal to the Cas9 cut site (3 bases upstream of PAM). Similarly, the forward primer for RH was selected as 20-24 bp immediately distal to the cut site. Keeping the LH reverse primer as constant (constraint), a forward primer was curated on the online tool.

Similarly, keeping forward primer of RH the constraint, a reverse primer for amplifying RH was curated with the same parameters as stated earlier. Further, to clone the arms with FRTs in the desired orientation and, in pBSKS via Takara® In-fusion (Cat# 638915), the forward primer was added with an overhang complementary to 15 bp immediately upstream of EcoRV digested pBSKS. The reverse primer was also incorporated with a 15 bp o/h homologous to 15 bp immediately downstream of EcoRV digested pBSKS. The reverse primer of LH was provided with an overhang for 2/3^rd^ the size of the 34 bp minimal FRT sequence:

5′-GAAGTTCCTATTCtctagaaaGtATAGGAACTTC-3′

Note that the nucleotides indicated by uppercase alphabets represent binding sites for subunits of the flippase enzyme (FLP). The lowercase alphabets indicate the asymmetric core of the FRT site, which gives the site a particular orientation. A similar o/h was also provided on the forward primer of RH, ensuring that it also has a 15 bps homology to the FRT overhang of LH reverse primer. The complete list of primers is provided in Supplemental Table S4-S6.

### Fly stocks and culture

Flies were grown in standard cornmeal yeast extract medium at 25°C unless otherwise specified.

### Generation of transgenic lines with FRT insertions at various regions within the Abd-B cis-regulatory landscape

All the donor constructs to target FRT were cloned in pBSKS vector with ∼1 Kb homology arms from the region flanking the site of FRT insertion. The arms were amplified with overhangs for FRT sequences with 15-20 bp homology. The homology arms with FRT sequences were cloned in EcoRV digested pBSKS vector using standard protocol for Takara® in-fusion assembly (Cat#102518). All the constructs to express the guideRNA were cloned in pCFD3 vectors as described earlier.

The donor and guide expressing constructs were co-injected in *nos-Cas9 Drosophila* embryos (BDSC#54591) as previously described or outsourced to Centre for Cellular and Molecular Platforms (C-CAMP), Bangalore, India. The injected G0s and G1 flies were crossed with third chromosome balancer as single fly crosses. The G1s were screened for the desired transgenes by using primers that eccentrically flanked the FRT site. The amplified product was digested using *Xba-I* to validate the presence of FRT prior to sequencing based confirmation. A detailed schematics for obtaining the FRT transgenes is shown in Supplemental Table S2-S3 and S7-S8.

### Generation of recombinants by Flp mediated recombination

The transgenic lines with desired FRTs were crossed and obtained with a cassette containing Flippase recombinase coding region under the influence of heat-shock inducible promoter (hsFLP; BDSC#7). The FLP activity was validated using a fly expressing Gal4 under the influence of Act5C promoter ubiquitously upon FRT excision (BDSC#4779) combined with UAS-GFP and hsFlp expressing flies (see supplementary information, Supplemental Fig. S2).

To obtain deletion and duplication, the flies with desired FRTs were crossed to obtain trans-heterozygous fly with respect to the position of FRT. The trans-heterozygous late second instar larvae are given heat shock at 37°C for 90 mins followed by a daily heat-shock periodically after every 24 hrs till eclosion. The detailed schematics of the crosses for obtaining deletion and duplication is shown in Supplemental Table S9-S10. For obtaining inversion, a fly with two FRTs in -cis is given heat shock as standardized. The schematics of genetic crosses are provided in Supplemental Table S11-S12.

### Population assay of variable phenotype for *Dp_i6-F8*

25 females and 15 males of either *Dp_i6-F8* or 25 females of *Dp_i6-F8* and 15 males of CS lines were mated and the number of progenies eclosing counted with different phenotypes. The distribution of heterozygous and homozygous progenies was recorded. The flies with affected abdominal segments A1-A8 were counted as category A. The flies with a distortion in A5-A8 were counted in category B. Category C included flies with aberrations in A6-A7. The flies having a defect in only one segment (often A6 or A7) were grouped as Category D. Category E mutants had minor defects in segments without perturbing the segmental boundaries, often characterized as wavy segmental borders. The category F flies appeared normal when compared to a wild-type fly. The distribution was normalized as the percentage of the total population for a given category. The detailed counting is presented in Supplementary Data in form of a workbook.

### Cuticle preparations of adult abdominal segments

Adult abdominal cuticles were prepared as described earlier. Briefly, 2-3 days old adult flies were dehydrated in 70% ethanol for 24 hrs. The flies are then boiled at 70°C in 10% potassium hydroxide for 90 mins followed by three washes of autoclaved distilled water. Next, the flies are washed in autoclaved water for 90 mins at 70°C and transferred to room temperature 70% ethanol.

### Mounting and imaging of cuticle preparations

The cuticles are transferred to a watch glass containing 70% ethanol and head-thorax region is separated from the abdominal segments by a dissection needle. The abdominal segments are given a sharp cut along the dorsal midline and transferred to a clean glass slide containing P700 halocarbon oil (Sigma #H8898). The cuticle is spread in desired orientation using dissection needles and covered with cover slip. The cuticles were imaged at 1.0X objective, 35X zoom on Zeiss AxioZoom V16 stereo microscope.

## Acknowledgments

Authors would like to acknowledge the members of RKM group for constant discussion during the conceptualization and execution of the project. NH is a fellow of Department of Biotechnology, Govt of India and thanks the agency for timely financial support. RKM is recipient of JC Bose National fellowship, India. The authors also thank CSIR, India, JC Bose National Fellowship, India, and Tata Institute for Genetics and Society, India for financial support at various levels. Further, the authors are thankful to Sonal Nagarkar Jaiswal lab for providing hsFlp and Act5C>STOP>Gal4; UAS-GFP flies, and, Centre for Cellular and Molecular Platforms (C-CAMP), Bangalore for injecting some of the constructs used in the study. The authors would especially like to acknowledge the arduous efforts of Mandar Naik, Binita Ghosh and Padmasini Chary in assisting NH with initial screenings of FRT transgenes and standardizations of heat shock experiments for the FLP expressing flies.

## Author Contributions

RKM conceptualized the project. RKM and NH designed the experiments. NH executed the project. SP assisted NH in a molecular screening of the recombinants, cuticle preparations, and imaging. NH wrote the manuscript with inputs from RKM. RKM supervised the project. NH and RKM edited the manuscript.

