## Supplementary Information for "CRISPR/Cas9 and FLP-FRT mediated multi-modular engineering of the *cis*-regulatory landscape of the bithorax complex of *Drosophila melanogaster*"

**Validation of FLP activity**

To generate the required genomic alterations, *Flp* was expressed under the influence of heat-shock inducible promoter (*hs*). the activation of FLP was validated prior to molecular screening using a transgenic line expressing *Gal4* under the control of *Act-5C* constitutively expressing promoter. Notedly, the promoter is followed by a 3XSTOP transcription termination site that is flanked by FRT sequences in same orientation as shown in Fig SF2. The fly also has a *UAS-GFP* insertion on another chromosome. The fly is genetically crossed with *hsFlp* expressing flies to bring the assay system in Flp background. In the absence of heat-shock, FLP is not produced as a result of which the Gal4 also fails to express. Further, upon heat shock, FLP acts on the FRTs that flank the 3XSTOP site downstream of *Act-5C* promoter (SF 2A). Since the two FRTs are in the same orientation, the 3XSTOP site is excised from the chromosome thereby allowing *Gal4* to express under *Act-5C* promoter constitutively. The flies carrying active flippase, thus, express GFP ubiquitously. Non-heat-shock controls and FRT lines without Flp did not express any GFP confirming the activity of Flp (SF 2B).

Note that the embryonic heat shocks were lethal to the animals wherein we altered the *cis-*regulatory landscape. This could be due to the primary role of *Abd-B* in embryonic development. Further, we could not obtain any recombinant when the heat-shock was given to the adults. We reasoned that a bulk of germ cells are already made in the flies leading to a pool of healthy haploids when the heat shock was given. Since, a majority of our flies required recombination across chromosomes, the adults would keep producing healthy embryos for a long time. Hence heat shock was given to late second instar larvae followed by periodic heat shocks at 24 hr interval till eclosion (SF 2C). The female gametogenesis begins in late second instar larval stages while the male gametogenesis initiates around late third instar stage. To establish a stable transgene and not just somatic mutations with clonal population, heat shocks were given in anticipation of required changes in the gametes. Later, the progenies were screened as described in methods.
