## Supplementary Material for "CRISPR/Cas9 and FLP-FRT mediated multi-modular engineering of the *cis*-regulatory landscape of the bithorax complex of *Drosophila melanogaster*"

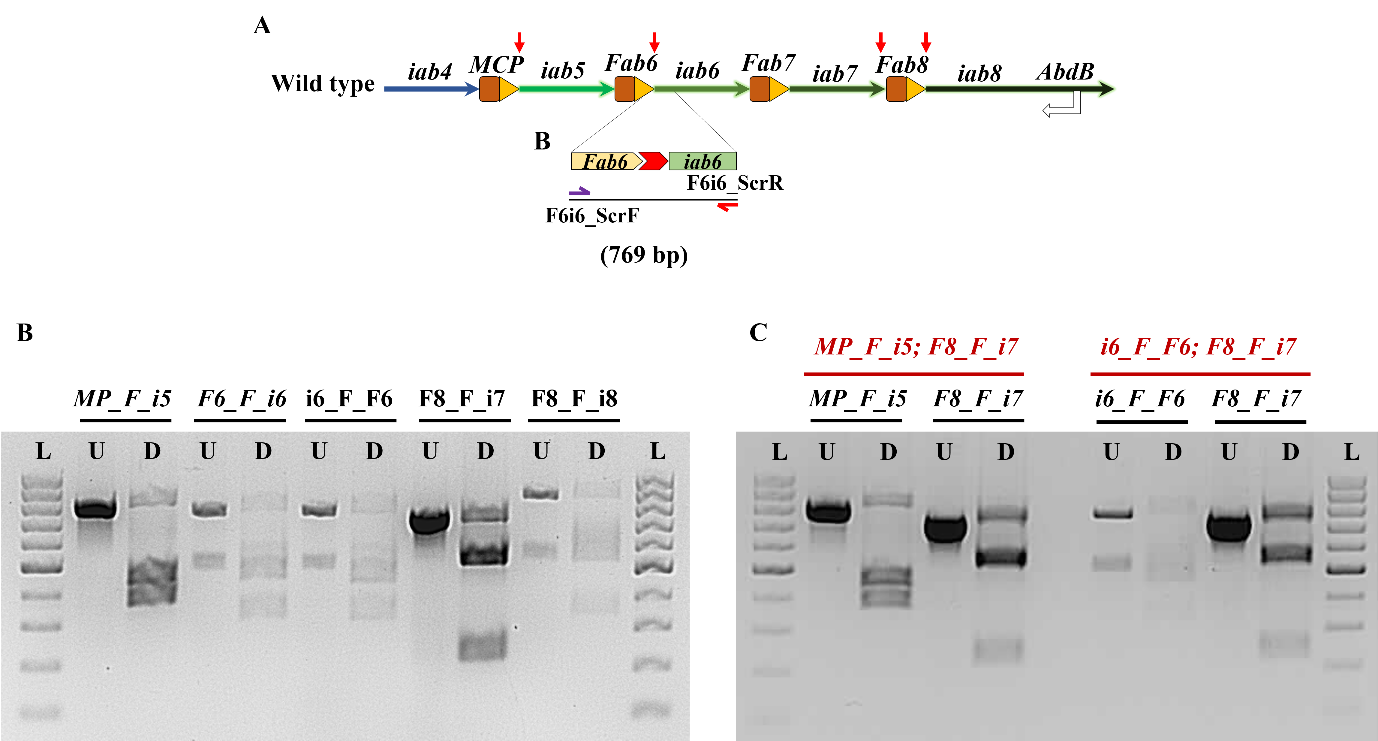

**Supplemental Figure S1: Molecular validation of FRT *knock-in*s at various regions within the bithorax complex:** (A) Representation of wild type locus exemplifying validation of FRT at *Fab6-iab6* locus. (B) Molecular validation of FRT at single locus. The genomic DNA of the transgenes is amplified with specific primers designed for the locus. Half of the amplicon is digested with *Xba-I* as a separate reaction and are run on the gel next to each other (U – undigested, D – digested; see text for details). (C) Molecular validation of double-FRT lines. The genomic DNA was amplified for multiple loci of interest and digested with *Xba-I*. The undigested and digested products are analysed in 1.5% agarose gel via electrophoresis.

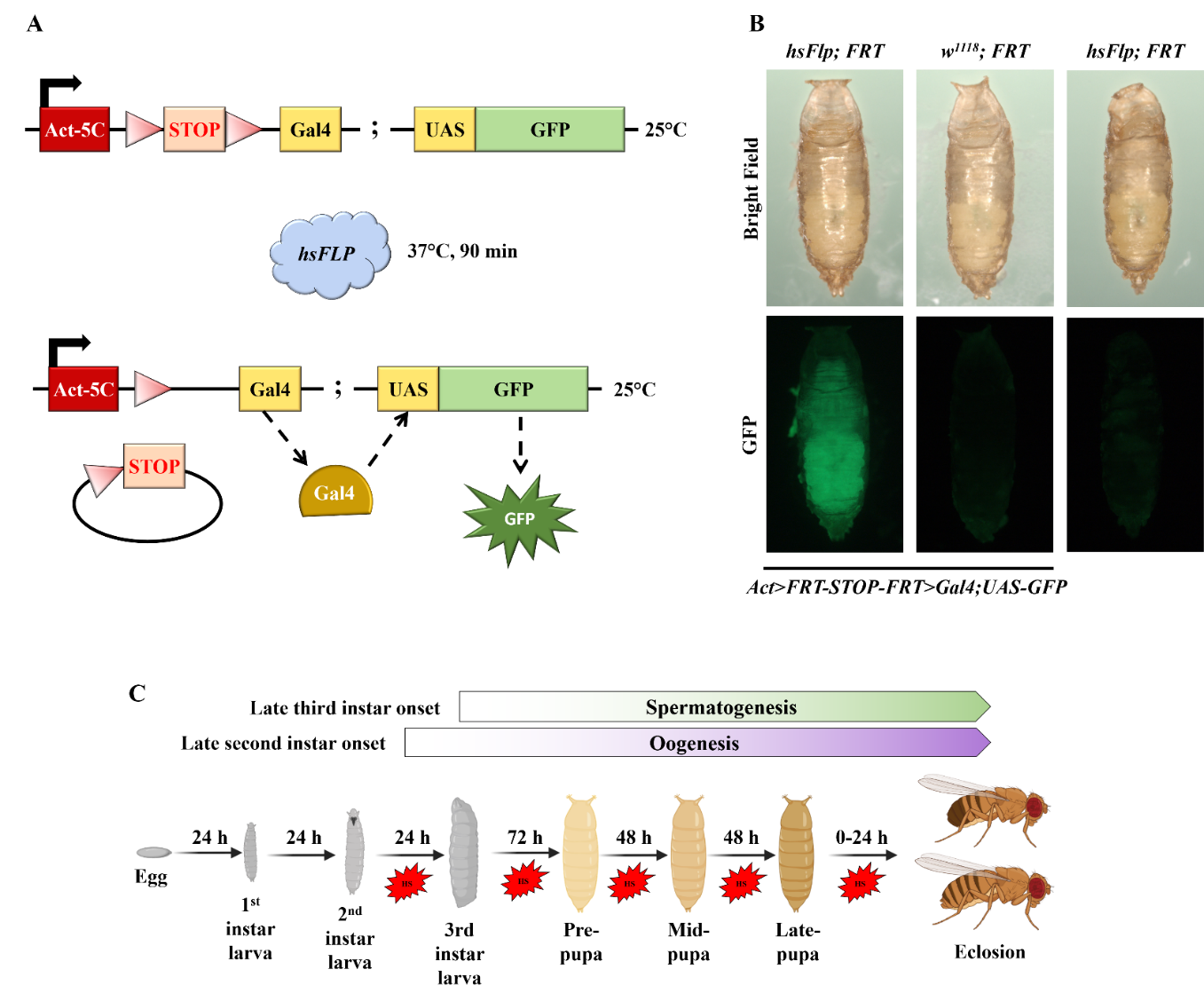

**Supplemental Figure S2: Validation of FLP activity:** (A) Representation of transgene used for validating activity of Flp. (B) All the flies involved in validating FLP activity are given heat shock at 37°C for 90 mins. Flies expressing functional flippase fluoresce green only when they have the hsFlp and Act-Gal4 module along with UAS-GFP. For flies with *w^1118^* and flies lacking Act5C-FRT.Gal4, no GFP is observed. (C) Strategy for providing heat shock to G1 animals from the late second instar stage. See text for details. Image created with [biorender](https://biorender.com/).

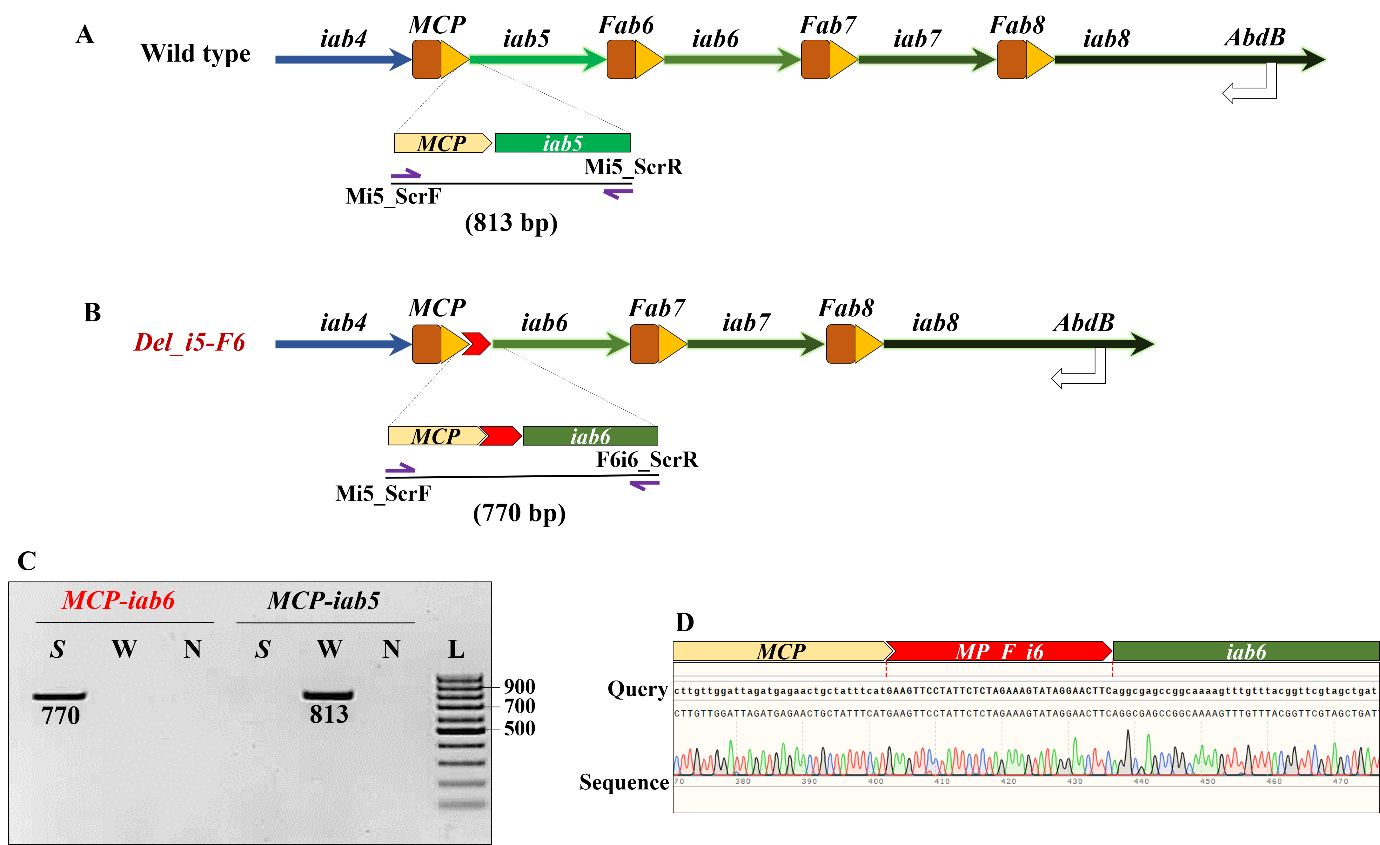

**Supplemental Figure S3: Molecular validation of the deletion of *iab5* through *Fab6*:** (A) A representation of wild-type *Abd-B* locus. The *MCP-iab5* region was selected as the control locus for PCR-based molecular validation. (B) After deletion, the region spanning *iab5* and *Fab6* was lost. However, due to FRT based recombination, the sites for recombination were specific. Therefore, the forward primer to screen FRT at *MCP-iab5* locus (Mi5_ScrF) was repurposed with the reverse primer to screen FRT insertion between *Fab6* and *iab6* (F6i6_ScrR). (C) 1.5% agarose gel with 1μg/ml EtBr shows amplification of specific products upon PCR. See text for details. The genomic DNA from the deletion mutant amplifies the recombined locus but fails to amplify the endogenous *MCP-iab5* locus as opposed to a wild-type CS fly. (D) Confirmation of deletion locus by sequencing. The region was sequenced using multiple primers.

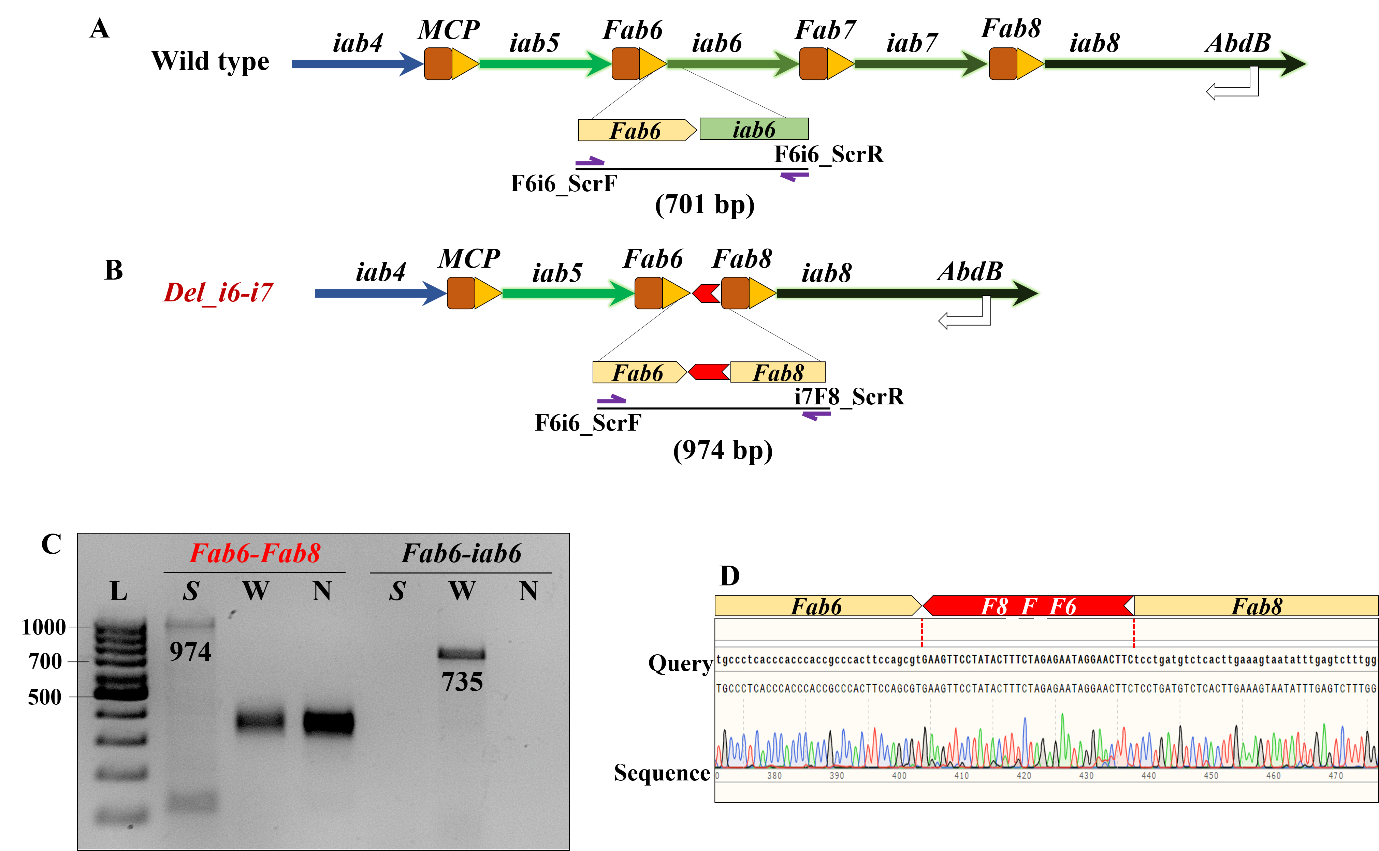

**Supplemental Figure S4: Molecular validation of the deletion of *iab6* through *iab7*:** (A) A representation of wild-type *Abd-B* locus. The region used as a PCR control is also depicted (outset from *Fab6-iab6* region). (B) After deletion, the region spanning *iab6* through *iab7* was lost. However, due to FRT based recombination, the sites for recombination are specific. Therefore, the forward primer to screen FRT at *Fab6-iab6* locus (F6i6_ScrF) was repurposed with the reverse primer to screen FRT insertion between *iab7* and *Fab8* (i7F8_ScrR). (C) 1.5% agarose gel with 1μg/ml EtBr shows amplification of specific products upon PCR. See text for details. The genomic DNA from the deletion mutant amplifies the recombined locus but fails to amplify the endogenous *Fab6-iab6* locus as opposed to a wild-type CS fly. (D) Confirmation of deletion locus by sequencing. The region was sequenced using multiple primers.

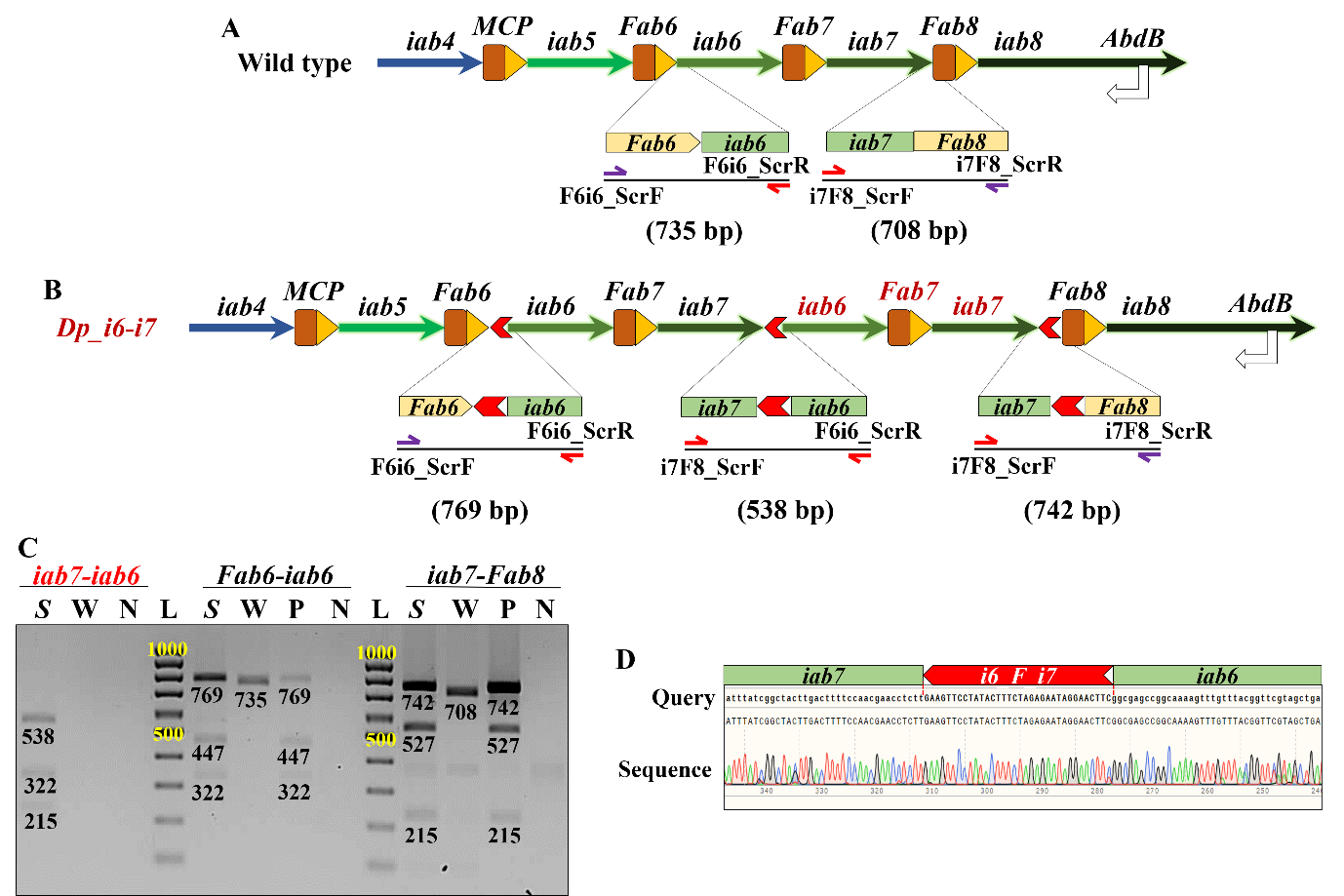

**Supplemental Figure S5: Molecular validation of the duplication of *iab6* through *iab7*:** (A) A representation of wild-type *Abd-B* locus. The regions used as a PCR control are also depicted (outset from *Fab6-iab6* and *iab7-Fab8*). (B) After duplication, the region spanning *iab6* through *iab7* is duplicated and juxtaposed next to the proximal *iab7*. The flies also have three FRTs as indicated in outsets representing loci amplified from the *Dp_i6-i7*. The forward primer to screen FRT at *iab7-Fab8* locus (i7F8_ScrF) was repurposed with the reverse primer to screen FRT insertion between *Fab6* and *iab6* (F6i6_ScrR). Only a fly with the desired duplication can amplify the genomic DNA with mentioned primers, thereby indicating the recombination event. (C) 1.5% agarose gel with 1μg/ml EtBr shows amplification of specific products upon PCR. The genomic DNA from the duplication mutant amplifies the recombined locus, which is not amplified by the genomic DNA of a CS fly. The *Fab6-iab6* and *iab7-Fab8* loci also amplify in the mutant indicating a duplication event. Further, the amplified product undergoes *Xba-I* mediated digestion from the duplication mutant (*Fab6-iab6*, *iab7-iab6* and *iab7-Fab8*). The amplified product from the CS fly from *Fab6-iab6,* and *iab7-Fab8* do not undergo restriction digestion via *Xba-I*. S – Sample, W – Wild-type, P – Positive control N- No template control. (D) Confirmation of duplication locus by sequencing.

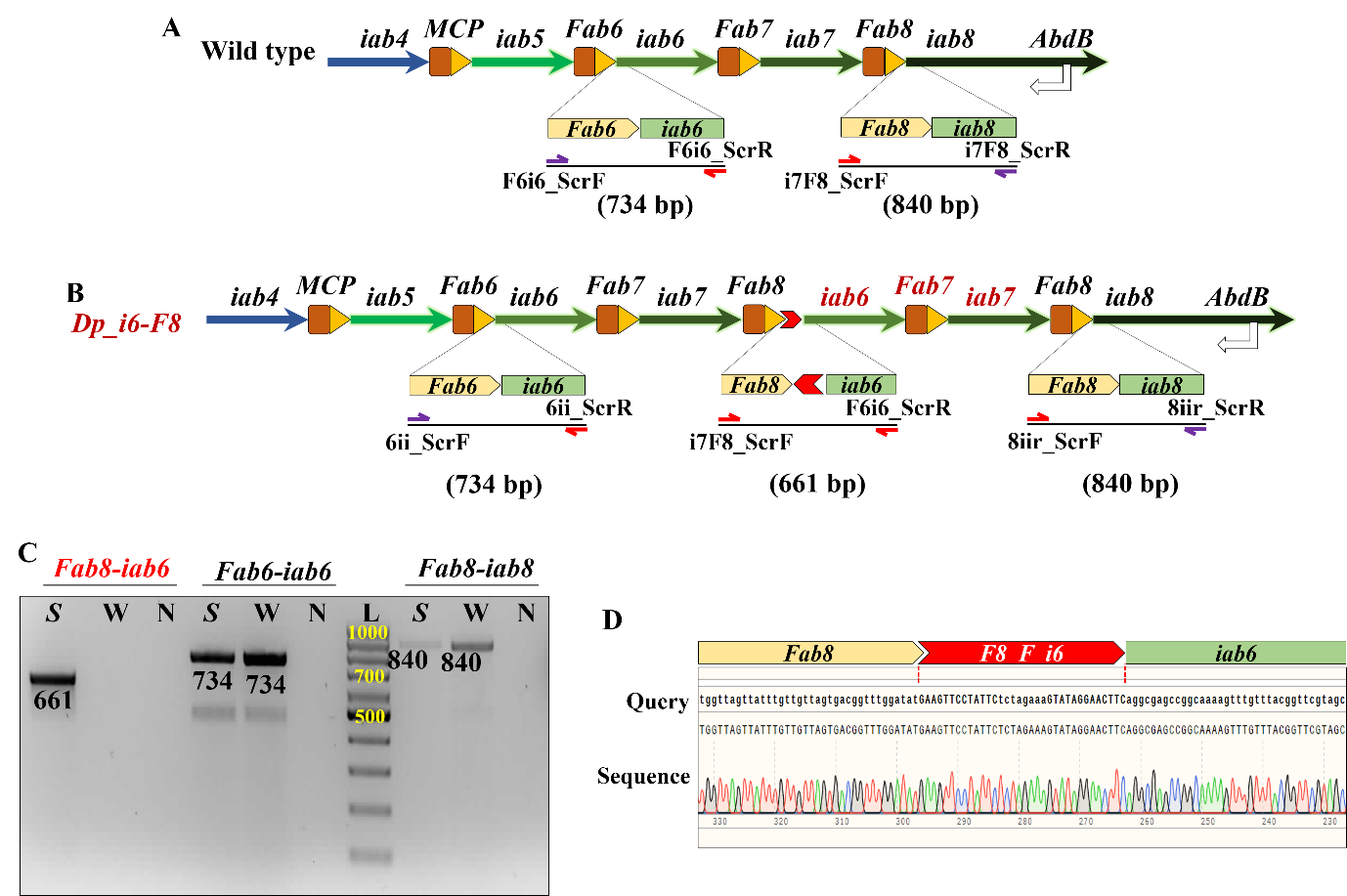

**Supplemental Figure S6: Molecular validation of the duplication of *iab6* through *Fab8*:** (A) A representation of wild-type *Abd-B* locus. The regions used as a PCR control is also depicted (outset from *Fab6-iab6* and *Fab8-iab8*). (B) After duplication, the region spanning *iab6* through *iab8* is duplicated and juxtaposed next to the proximal *Fab8*. The forward primer to screen FRT at *Fab8-iab8* locus (F8i8_ScrF) was repurposed with the reverse primer to screen FRT insertion between *Fab6* and *iab6* (F6i6_ScrR). Only a fly with the desired duplication can amplify the genomic DNA with mentioned primers, thereby indicating the recombination event. (C) 1.5% agarose gel with 1μg/ml EtBr shows amplification of specific products upon PCR. The genomic DNA from the duplication mutant amplifies the recombined locus, which is not amplified by the genomic DNA of a CS fly. The *Fab6-iab6* and *Fab8-iab8* loci amplify in the mutant and the CS fly, indicating a duplication event. (D) Confirmation of duplication locus by sequencing.

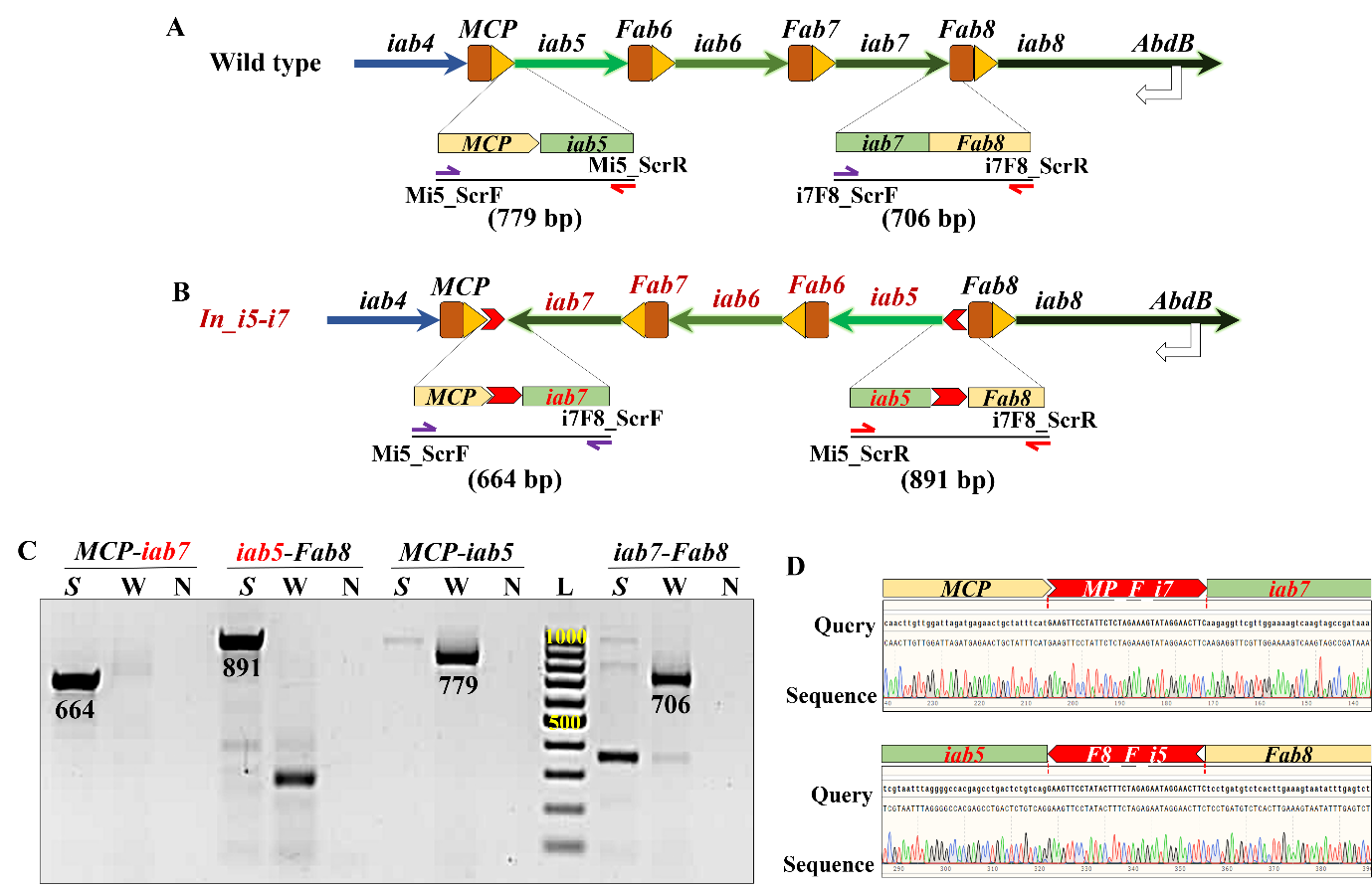

**Supplemental Figure S7: Molecular validation of the inversion of *iab5* through *iab7*:** (A) A representation of wild-type *Abd-B* locus. The regions used as a PCR control is also depicted (outset from *MCP-iab5* and *iab7-Fab8*). (B) After inversion, the region spanning *iab5* through *iab7* is inverted. Due to FRT based recombination, the sites for recombination are specific as indicated in outsets representing the loci amplified from the *In_i5-i7* flies (*MCP-iab7* and *iab5-Fab8*). The forward primer to screen FRT at *MCP-iab5* locus (Mi5_ScrF) was repurposed with the forward primer to screen FRT insertion between *iab7* and *Fab8* (i7F8_ScrF) to screen *MCP-iab7*. A second locus of recombination (*iab5-Fab8*) was amplified by repurposing the reverse primers for screening the *MCP-iab5* (Mi5_ScrR) and *iab7-Fab8* (i7F8_ScrR) loci in the wild-type flies. Only a fly with the desired inversion can amplify the genomic DNA with mentioned primers. (C) 1.5% agarose gel with 1μg/ml EtBr shows amplification of specific products upon PCR. See text for details. The genomic DNA from the inversion mutant amplifies the recombined locus, which is not amplified by the genomic DNA of a CS fly. On the contrary, the *MCP-iab5* and *iab7-Fab8* loci amplify in the CS fly, not in the inversion mutant. (D) Confirmation of inversion by Sanger sequencing.

| **S. No.** | **Name** | **Genotype** | **Site of insertion of FRT or regions affected upon recombination*** |
| --- | --- | --- | --- |
| 1 | *MP_F_i5* | *Abd-B^[MCP-KI{FRT}-iab5]^* | 16,872,061 |
| 2 | *F6_F_i6* | *Abd-B^[Fab6-KI{FRT}-iab6]^* | 16,887,207 |
| 3 | *i6_F_F6* | *Abd-B^[iab6-KI{FRT}-Fab6]^* | 16,887,207 |
| 4 | *F8_F_i7* | *Abd-B^[Fab8-KI{FRT}-iab7]^* | 16,918,581 |
| 5 | *F8_F_i8* | *Abd-B^[Fab8-KI{FRT}-iab8]^* | 16,921,983 |
| 6 | *i6_F_F6, F8_F_i7* | *Abd-B^[iab6-KI{FRT}-Fab6, Fab8-KI{FRT}-iab7]^* | *i6_F_F6* at 16,887,207  *F8_F_i7* at 16,918,581 |
| 7 | *MP_F_i5, F8_F_i7* | *Abd-B^[MCP-KI{FRT}-iab5, Fab8-KI{FRT}-iab7]^* | *i6_F_F6* at 16,887,207  *F8_F_i7* at 16,918,581 |
| 8 | *Del_i5-F6* | *Abd-B^[Del(iab5-Fab6)]^* | 16,872,062..16,887,207 |
| 9 | *Del_i6-i7* | *Abd-B^[Del(iab6-iab7)]^* | 16,887,208..16,918,581 |
| 10 | *Dp_i6-i7* | *Abd-B^[Dp(iab6-iab7)]^* | 16,887,208..16,918,581 |
| 11 | *Dp_i6_F8* | *Abd-B^[Dp(iab6-Fab8)]^* | 16,887,208..16,921,983 |
| 12 | *Inv_i5-i7* | *Abd-B^[Inv(iab5-iab7)]^* | 16,872,062.. 16,918,581 |

* All coordinates are mentioned as per the latest *Drosophila melanogaster* genome assembly coordinates R6.0

**Supplemental Table S1: List of transgenic lines and FRT mediated recombinants generated in the study:** The FRT sequences are inserted in *Drosophila* genome between *MCP-iab5*, *Fab6-iab6*, *iab7-Fab8,* and *Fab8-iab8*. The nomenclature is displayed with respect to the orientation of FRT sequence. For example, line *MP_F_i5* has FRT inserted between *MCP* and *iab5* with the direction of FRT towards *iab5*. The table also lists different mutations generated as a result of Flp-FRT mediated recombination.

| **S. No.** | **Name** | **Purpose** |
| --- | --- | --- |
| 1 | pMP_F_i4 | Donor to insert FRT between *MCP* and *iab4* towards *iab4* |
| 2 | pMP_F_i5 | Donor to insert FRT between *MCP* and *iab5* towards *iab5* |
| 3 | pi5_F_MP | Donor to insert FRT between *iab5* and *MCP* towards *MCP* |
| 4 | pF6_F_i6 | Donor to insert FRT between *Fab6* and *iab6* towards *iab6* |
| 5 | pi6_F_F6 | Donor to insert FRT between *iab6* and *Fab6* towards *Fab6* |
| 6 | pF7_F_i6 | Donor to insert FRT between *Fab7* and *iab6* towards *iab6* |
| 7 | pF7_F_i7 | Donor to insert FRT between *Fab7* and *iab7* towards *iab7* |
| 8 | pF8_F_i7 | Donor to insert FRT between *Fab8* and *iab7* towards *iab7* |
| 9 | pF8_F_i8 | Donor to insert FRT between *Fab8* and *iab8* towards *iab8* |
| 10 | pi8_F_F8 | Donor to insert FRT between *iab8* and *Fab8* towards *Fab8* |

**Supplemental Table S2: List of donor constructs generated in pBSKS(-) for targeting FRT via CRISPR/Cas9:** The FRT sequences were cloned in different orientations with required homology arms in pBSKS(-) vector backbone. See methods for details.

| **S. No.** | **Name** | **Purpose** |
| --- | --- | --- |
| 1 | pi4-MP_C | Guide to target region between *iab4* and *MCP* |
| 2 | pMP-i5_C | Guide to target region between *MCP* and *iab5* |
| 3 | pi5-F6_C | Guide to target region between *iab5*and *Fab6* |
| 4 | pF6-i6_C | Guide to target region between *Fab6* and *iab6* |
| 5 | pi6-F7_C | Guide to target region between *iab6* and *Fab7* |
| 6 | pF7-i7_C | Guide to target region between *Fab7* and *iab7* |
| 7 | pi7-F8_C | Guide to target region between *iab7* and *Fab8* |
| 8 | pF8-i8_C | Guide to target region between *Fab8* and *iab8* |

**Supplemental Table S3: List of guide RNA expressing constructs generated in pCFD3 vector for targeting FRT:** Different guides were selected and cloned in pCFD3 vector. See methods for details.

| **S. No.** | **Name** | **Sequence (5’ → 3’)** | **Purpose** |
| --- | --- | --- | --- |
| 1 | pbsksEcoRV_LH_i4MP | Ctgcaggaattcgatgcttactgggtaaagcggat | Amplify left homology arm with left half of the FRT for the Gibson® assembly cloning of pMP_F_i4 |
| 2 | i4MP_LH_LFRT | ttctctagaaagtataggaacttcacagcatttcaggtgccacgc |  |
| 3 | i4MP_RH_RFRT | atactttctagagaataggaacttctaaaagccagctgcagactt | Amplify right homology arm with right half of the FRT for the Gibson® assembly cloning of pMP_F_i4 |
| 4 | RH_i4MP_pbsksEcoRV | atcgataagcttgatccgcattaaatattgcatgc |  |
| 5 | pbsksEcoRV_LH_MPi5 | ctgcaggaattcgatatacagcttccggcagcgac | Amplify left homology arm with left half of the FRT for the Gibson® assembly cloning of pMP_F_i5 |
| 6 | MPi5_LH_LFRT | atactttctagagaataggaacttctgaaatagcagttctcatc |  |
| 7 | MPi5_RH_RFRT | ttctctagaaagtataggaacttcactgacagagtcaggctcgtg | Amplify right homology arm with right half of the FRT for the Gibson® assembly cloning of pMP_F_i5 |
| 8 | RH_MPi5_pbsksEcoRV | atcgataagcttgatagccactgataaattaccag |  |
| 9 | i5MP_LH_LFRT | ttctctagaaagtataggaacttcatgaaatagcagttctcatct | Can be repurposed with Primer# 5 and 8 respectively for cloning pi5_F_MP |
| 10 | i5MP_RH_RFRT | atactttctagagaataggaacttcctgacagagtcaggctcgtg |  |
| 11 | pbsksEcoRV_LH_F6i6 | ctgcaggaattcgatcaaataaaccagcgctgcaa | Amplify left homology arm with left half of the FRT for the Gibson® assembly cloning of pF6_F_i6 |
| 12 | F6i6_LH_LFRT | atactttctagagaataggaacttccgctggaagtgggcggtggg |  |
| 13 | F6i6_RH_RFRT | ttctctagaaagtataggaacttcaggcgagccggcaaaagtttg | Amplify right homology arm with right half of the FRT for the Gibson® assembly cloning of pF6_F_i6 |
| 14 | RH_F6i6_pbsksEcoRV | atcgataagcttgattgtgttatttggcgaaacag |  |
| 15 | i6F6_LH_LFRT | ttctctagaaagtataggaacttcacgctggaagtgggcggtggg | Can be repurposed with Primer# 11 and 14 respectively for cloning pi6_F_F6 |
| 16 | i6F6_RH_RFRT | atactttctagagaataggaacttcggcgagccggcaaaagtttg |  |
| 17 | pbsksEcoRV_LH_i6F7 | ctgcaggaattcgataggcgacgctttgtcataaa | Amplify left homology arm with left half of the FRT for the Gibson® assembly cloning of pF7_F_i6 |
| 18 | i6F7_LH_LFRT | ttctctagaaagtataggaacttcagagaaattacttttctgtga |  |
| 19 | i6F7_RH_RFRT | atactttctagagaataggaacttccaactcgtgaaggtcgtgca | Amplify right homology arm with right half of the FRT for the Gibson® assembly cloning of pF7_F_i6 |
| 20 | RH_i6F7_pbsksEcoRV | atcgataagcttgattgagcgacattctacctcgc |  |
| 21 | pbsksEcoRV_LH_F7i7 | ctgcaggaattcgatcaccgcaaaatccaattgga | Amplify left homology arm with left half of the FRT for the Gibson® assembly cloning of pF6_F_i6 |
| 22 | F7i7_LH_LFRT | atactttctagagaataggaacttcggtctacaaaggtctattta |  |
| 23 | F7i7_RH_RFRT | ttctctagaaagtataggaacttcatcttaccgaacccaataatt | Amplify right homology arm with right half of the FRT for the Gibson® assembly cloning of pF6_F_i6 |
| 24 | RH_F7i7_pbsksEcoRV | atcgataagcttgattttagcagcggaacggtagt |  |
| 25 | pbsksEcoRV_LH_i7F8 | ctgcaggaattcgatcatgccgccattcggcgatt | Amplify left homology arm with left half of the FRT for the Gibson® assembly cloning of pF8_F_i7 |
| 26 | i7F8_LH_LFRT | ttctctagaaagtataggaacttcaagaggttcgttggaaaagtc |  |
| 27 | i7F8_RH_RFRT | atactttctagagaataggaacttctcctgatgtctcacttgaaa | Amplify right homology arm with right half of the FRT for the Gibson® assembly cloning of pF8_F_i7 |
| 28 | RH_i7F8_pbsksEcoRV | atcgataagcttgatactcttgttatcaaagcaga |  |
| 29 | pbsksEcoRV_LH_F8i8 | ctgcaggaattcgatgcaacatgtggcattaatta | Amplify left homology arm with left half of the FRT for the Gibson® assembly cloning of pF8_F_i8 |
| 30 | F8i8_LH_LFRT | atactttctagagaataggaacttccacggtgctttggatgcgga |  |
| 31 | F8i8_RH_RFRT | ttctctagaaagtataggaacttcacacggtgctttggatgcgga | Amplify right homology arm with right half of the FRT for the Gibson® assembly cloning of pF8_F_i8 |
| 32 | RH_F8i8_pbsksEcoRV | atcgataagcttgatacggcctttgtcttcggtgg |  |
| 33 | i8F8_LH_LFRT | ttctctagaaagtataggaacttcaatatccaaaccgtcactaac | Can be repurposed with Primer# 29 and 32 respectively for cloning pi8_F_F8 |
| 34 | i8F8_RH_RFRT | atactttctagagaataggaacttcatatccaaaccgtcactaac |  |

**Supplemental Table S4: List of oligos for cloning donor constructs with FRTs in pBSKS:** The oligos were used for amplifying different homology arms with a particular orientation of FRT sequence. See methods for details.

| **S. No.** | **Name** | **Sequence** | **Target region for Cas9 mediated DSB** |
| --- | --- | --- | --- |
| 1 | i4MP_Cr_s | gtcggctggcttttacagcatttc | Between *iab4* and *MCP* |
| 2 | i4MP_Cr_as | aaacgaaatgctgtaaaagccagc |  |
| 3 | MPi5_Cr_s | gtcggctatttcactgacagagtc | Between *MCP* and *iab5* |
| 4 | MPi5_Cr_as | aaacgactctgtcagtgaaatagc |  |
| 5 | F6i6_Cr_s | gtcggccggctcgcccgctggaag | Between *Fab6* and *iab6* |
| 6 | F6i6_Cr_as | aaaccttccagcgggcgagccggc |  |
| 7 | i6F7_Cr_s | gtcggtaatttctccaactcgtga | Between *iab6* and *Fab7* |
| 8 | i6F7_Cr_as | aaactcacgagttggagaaattac |  |
| 9 | F7i7_Cr_s | gtcggttcggtaagaggtctacaa | Between *Fab7* and *iab7* |
| 10 | F7i7_Cr_as | aaacttgtagacctcttaccgaac |  |
| 11 | i7F8_Cr_s | gtcggacatcaggaagaggttcgt | Between *iab7* and *Fab8* |
| 12 | i7F8_Cr_as | aaacacgaacctcttcctgatgtc |  |
| 13 | F8i8_Cr_s | gtcggtttggatatcacggtgctt | Between *Fab8* and *iab8* |
| 14 | F8i8_Cr_as | aaacaagcaccgtgatatccaaac |  |

**Supplemental Table S5: List of oligos for cloning sgRNA expressing element in *pCFD3*:** The primers were annealed and ligated in the pCFD3 vector to yield sgRNA expressing constructs. See methods for details.

| **S. No.** | **Name** | **Sequence** | **Molecular validation of the transgenic line** |
| --- | --- | --- | --- |
| 1 | Mi5_ScrF | ccaccgcacttttctgttcg | *MP_F_i5* and *i5_F_MP* |
| 2 | Mi5_ScrR | tcagccagttccaaggttcc |  |
| 3 | F6i6_ScrF | tgtggctgctacgtgagttt | *F6P_F_i6* and *i6_F_F6P* |
| 4 | F6i6_ScrR | ttacttaaacgggccagcgt |  |
| 5 | i7F7_ScrF | ttgcatatcttcgcagggct | *F7B_F_i6* |
| 6 | i7F7_ScrR | ttccattcccgattgccgtt |  |
| 7 | F7i7_ScrF | acggcattttgttgggcatt | *F7P_F_i7* |
| 8 | F7i7_ScrR | gcaagggaattgtgtggacg |  |
| 9 | i7F8_ScrF | gcatcatggcctcggttttc | *F8B_F_i7* |
| 10 | i7F8_ScrR | atgtgtttgtgctggttggc |  |
| 11 | F8i8_ScrF | tcttggccgcattttgcatc | *F8P_F_i8* and *i8_F_F8P* |
| 12 | F8i8_ScrR | gaacggaaaccgcctcaaag |  |

**Supplemental Table S6: List of oligos for screening FRT insertion at genomic DNA in transgenes:** Different primers were designed eccentric to the FRT site to yield products of different lengths upon *Xba-I* mediated restriction digestion. These primers were used to screen the insertion of FRT as well as were repurposed to screen Flp-FRT mediated recombination events.

| **S. No.** | **Name** | **Genotype** | **Comments** |
| --- | --- | --- | --- |
| 1 | nos-Cas9 | y^[1]^ M{w^[+mC]^=nos-Cas9.P}ZH-2A w^[*]^ | BDSC# 54591. The fly produces Cas9 protein driven by nanos promoter expressing in pole cells of the embryo which form the germinal epithelium of the adult flies |
| 2 | T3/T6B | w^[118]^; + ; TM3Sb/TM6BTb | Third chromosome balancer |
| 3 | TG | Transgene of interest |  |
| 4 | Bal1 | Any one type of balancer: TM3Sb or TM6BTb |  |

**Supplemental Table S7: Nomenclature of flies used for obtaining transgenes:** The flies and genetic crosses mentioned below were utilized to obtain *knock-in* transgenic flies for FRT.

| **Generation** |  | **Fly 1** | **Fly 2** | **Comments** |
| --- | --- | --- | --- | --- |
| G0 | Genotype | nos-Cas9 | T3/T6B | Desired constructs are injected in the embryos of Fly1 |
|  | Phenotype | Yellow body, red eyes | Stubble, tubby, humeral bristles, ebony |  |
| G1 | Genotype | TG/Bal | TG/T6B | After 3 days of setting up single-fly crosses, flies are confirmed via *PC*R |
|  | Phenotype | Any of the balancer phenotype | Stubble, tubby, humeral bristles, ebony |  |
| G2 | Genotype | TG/Bal1 | TG/Bal1 | These crosses are set only for the confirmed G1 lines |
|  | Phenotype | Flies with same balancer are crossed | |  |
| G3 | Genotype | TG/TG | TG/TG | These flies are maintained as stocks |
|  | Phenotype | Normal body | |  |

**Supplemental Table S8: Genetic crosses for obtaining transgenic FRT *knock-in* flies**

| **S. No.** | **Name** | **Genotype** | **Comments** |
| --- | --- | --- | --- |
| 1 | *hsFlp; MP_F_i5* | P{ry[+t7.2]=hsFLP}1, y[1] w[1118]; MP_F_i5 | FRT inserted between *MCP*-PRE and *iab*5 with the orientation of FRT towards *iab*5 in hsFlp background |
| 2 | *hsFlp; F6_F_i6* | P{ry[+t7.2]=hsFLP}1, y[1] w[1118]; F6_F_i6 | FRT inserted between *Fab6* and *iab6* with the orientation of FRT towards *iab*6 in hsFlp background |
| 3 | T3/T6B | w^1118^; + ; TM3Sb/TM6BTb | Third chromosome balancer |
| 4 | #; *Del or Dup*i5_F6/Bal1* | #; +; *Del* or *Dupi5_F6*/Bal1 | **#** - X chromosome could be P{ry[+t7.2]=hsFLP}1, y[1] w[1118] or w[1118] depending on the parent and is immaterial to the cross  **Del* or Dup*** – Deletion or Duplication, to be validated molecularly |

**Supplemental Table S9: Nomenclature of flies used for FLP mediated recombination to obtain the deletion or duplication mutants:** If the two FRTs are in a trans-heterozygous position in the same orientation, the resultant recombination would lead to the generation of a chromosome with deletion while the other with duplication of the region that they flank. Therefore, we utilized the mentioned flies to obtain the desired manipulation. A case for Deletion/Duplication of *iab5-Fab6* is exampled in the table.

| **Generation** |  | **Fly line 1** | **Fly line 2** | **Comments** |
| --- | --- | --- | --- | --- |
| G0 | Genotype | *hsFlp; MP_F_i5* | *hsFlp; F6_F_i6* |  |
|  | Phenotype | Yellow body, white eyes | Yellow body, white eyes |  |
| G1 Larvae | Genotype | *hsFlp; MP_F_i5/F6_F_i6* | | The transheterozygous larvae were given heat shock at 37^o^C from late 2nd instar onwards to late pupal stages |
| G1 (adults eclosed) | Genotype | *hsFlp; MP_F_i5/F6_F_i6* | T3/T6B | Fly line 1 is eclosing after heat shock. |
|  | Phenotype | White eyes | White eyes, stubble bristles, humeral bristles, tubby, ebony body |  |
| G2^ | Genotype | #; *Del or Dup*i5_F6*/Bal1 | T3/T6B | # - genotype on X chromosome is immaterial  * - The chromosome could be a deletion, duplication or similar to any of G0 flies.  ^Pools of 10 flies each are set-up and screened after 3-4 days via PCR.  Positive pools are propagated via single-fly crosses |
|  | Phenotype | Any one balancer phenotype T3 or T6; might also have a homeotic phenotype | White eyes, stubble bristles, humeral bristles, tubby, ebony body |  |
| G3 | Genotype | #; *Del or Dup*i5_F6*/Bal1 | T3/T6B | # and * same as above.  Single fly crosses are set from the pool that was positive for a transgene |
|  | Phenotype | Any one balancer phenotype T3 or T6; might also have a homeotic phenotype | White eyes, stubble bristles, humeral bristles, tubby, ebony body |  |
| G4 | Genotype | #; *Del or Dup*i5_F6*/Bal1 | #; Del or Dup*/Bal1 | * These are the confirmed deletion or duplication lines via *PC*R from G3 single fly cross |
|  | Phenotype | Any one balancer phenotype T3 or T6; might also have a homeotic phenotype | Any one balancer phenotype T3 or T6; might also have a homeotic phenotype |  |
| G5 | Genotype | #; *Del or Dup*i5_F6*/Del or Dup*i5_F6 | #; Del or Dup*i5_F6/Del or Dup*i5_F6 | *Stock of confirmed deletion or duplication line |
|  | Phenotype | White eyes, putative homeotic transformation | |  |

**Supplemental Table S10: Genetic crosses to obtain a deletion or a duplication mutant:** If the two FRTs are in a trans-heterozygous position in the same orientation, the resultant recombination would lead to the generation of a chromosome with deletion while the other with duplication of the region that they flank. A case for Deletion/Duplication of *iab5-Fab6* is exampled in the table. Deletions and duplications of other regions were generated in a similar manner. These regions include the deletion or duplication of *iab6-iab7* obtained from *i6_F_F6*, *F8_F_i7* double FRT flies, and, deletion or duplication of *iab6-Fab8* obtained from *F6_F_i6* and *F8_F_i8*.

| **S. No.** | **Name** | **Genotype** | **Comments** |
| --- | --- | --- | --- |
| 1 | hsFlp; MP_F_i5, F8_F_i7 | P{ry[+t7.2]=hsFLP}1, y[1] w[1118]; MP_F_i5, F8_F_i7 | FRT1 inserted between *MCP*-PRE and *iab*5 with the orientation of FRT towards *iab*5 while the FRT2 is inserted between *Fab*8-Boundary and *iab*7 with the orientation towards *iab*7 in hsFlp background. Thus, the two FRTs face each otheron the same chromosome. |
| 3 | T3/T6B | w118; + ; TM3Sb/TM6BTb | Third chromosome balancer |
| 4 | #; In*i5_i7/Bal1 | #; +; In_i5-i7/Bal1 | # - X chromosome could be P{ry[+t7.2]=hsFLP}1, y[1] w[1118] or w[1118] depending on the parent and is immaterial to the cross  **In*** – Inversion to be screened molecularly |

**Supplemental Table S11: Nomenclature of flies used for generating FLP mediated inversion of *iab5-iab7*:** Inversions are obtained when the two FRTs are in the opposite orientation and in *-cis*. Therefore, we utilized the listed flies to obtain the desired manipulation.

| **Generation** |  | **Fly line 1** | **Fly line 2** | **Comments** |
| --- | --- | --- | --- | --- |
| G0 | Genotype | hsFlp; MP_F_i5, F8_F_i7 | T3/T6B |  |
|  | Phenotype | Yellow body, white eyes | Yellow body, white eyes |  |
| G1 Larvae | Genotype | hsFlp; MP_F_i5, F8_F_i7/ Bal1 | T3/T6B | The heterozgous larvae were given heat shock at 37oC from late 2nd instar onwards to late pupal stages |
| G1 (adults eclosed) | Genotype | hsFLP; MP_F_i5, F8_F_i7/Bal1 | T3/T6B | Fly line 1 is eclosing after heat shock. |
|  | Phenotype | Stubble bristles, white eyes | White eyes, stubble bristles, humeral bristles, tubby, ebony body |  |
| G2 | Genotype | #; In*i5_i7/Bal1 | T3/T6B | # - genotype on X chromosome is immaterial, * - The chromsome is a potential inversion of the desired region or can be similar to the G0 fly. Pools of 10 flies each are set-up and screened after 3-4 days via *PC*R |
|  | Phenotype | Any one balancer phenotype T3 or T6; might also have a homeotic phenotype | White eyes, stubble bristles, humeral bristles, tubby, ebony body |  |
| G3 | Genotype | #; In*i5_i7/Bal1 | T3/T6B | # and * same as above. Single fly crosses are set from the pool that was positive for a transgene |
|  | Phenotype | Any one balancer phenotype T3 or T6; might also have a homeotic phenotype | White eyes, stubble bristles, humeral bristles, tubby, ebony body |  |
| G4 | Genotype | #; In*i5_i7/Bal1 | #; In*i5_i7/Bal1 | * These are the confirmed inversion lines via *PC*R for G3 single fly cross |
|  | Phenotype | Any one balancer phenotype T3 or T6; might also have a homeotic phenotype | Any one balancer phenotype T3 or T6; might also have a homeotic phenotype |  |
| G5 | Genotype | #; In-i5_i7/Bal1 | #; In-i5_i7/Bal1 | Stock of confirmed inversion line |
|  | Phenotype | White eyes, putative homeotic transformation | |  |

**Supplemental Table S12: Genetic crosses to obtain a mutant with an inversion of *iab5-iab7***
